# Defective cyclophilin A induces TDP-43 proteinopathy: implications for amyotrophic lateral sclerosis and frontotemporal dementia

**DOI:** 10.1101/2020.06.08.129528

**Authors:** Laura Pasetto, Maurizio Grassano, Silvia Pozzi, Silvia Luotti, Eliana Sammali, Alice Migazzi, Manuela Basso, Giovanni Spagnolli, Emiliano Biasini, Edoardo Micotti, Milica Cerovic, Mirjana Carli, Gianluigi Forloni, Giovanni De Marco, Umberto Manera, Cristina Moglia, Gabriele Mora, Bryan J. Traynor, Adriano Chiò, Andrea Calvo, Valentina Bonetto

## Abstract

Aggregation and cytoplasmic mislocalization of TDP-43 are pathological hallmarks of amyotrophic lateral sclerosis (ALS) and frontotemporal dementia (FTD) spectrum. However, the molecular mechanism by which TDP-43 aggregates form and cause neurodegeneration remains poorly understood. Cyclophilin A, also known as peptidyl-prolyl *cis-trans* isomerase A (PPIA), is a foldase and molecular chaperone. We previously found that PPIA interacts with TDP-43 and governs some of its functions, and its deficiency accelerates disease in a mouse model of ALS. Here we characterized *PPIA* knock-out mice throughout their lifespan and found that they develop a neurodegenerative disease with key behavioural features of FTD, marked TDP-43 pathology and late-onset motor dysfunction. In the mouse brain, deficient PPIA induces aggregation of the GTP-binding nuclear protein Ran, a PPIA substrate required for TDP-43 nucleocytoplasmic trafficking. Moreover, in absence of PPIA, TDP-43 autoregulation is perturbed and TDP-43 and proteins involved in synaptic function are downregulated, leading to impairment of synaptic plasticity. Finally, we found that PPIA was downregulated in several ALS and ALS-FTD patients and identified a PPIA loss-of-function mutation in a sporadic ALS patient. The mutant PPIA has low stability, altered structure and impaired interaction with TDP-43. These findings strongly implicate that defective PPIA function causes TDP-43 mislocalization and dysfunction and should be considered in future therapeutic approaches.

## Introduction

TAR DNA-binding protein-43 (TDP-43), encoded by the *TARDBP* gene, is a predominantly nuclear RNA/DNA-binding protein that exerts vital functions in various steps of RNA metabolism, shuttling between the nucleus and the cytoplasm^1^. To do that TDP-43 physiological levels and correct localization have to be tightly controlled also through autoregulation^2, 3^.

Cytoplasmic mislocalization of TDP-43 and aggregation of its hyperphosphorylated, ubiquitinated and C-terminal fragmented forms are common histopathological hallmarks of the amyotrophic lateral sclerosis (ALS) and frontotemporal dementia (FTD) disease spectrum^4, 5^. TDP-43 proteinopathy is observed in ∼97% of ALS and ∼50% of FTD patients, in brain and spinal cord, but also in peripheral blood mononuclear cells (PBMCs)^6, 7^. *TARDBP* mutations in ALS and FTD cases have established a direct link between TDP-43 abnormalities and neurodegeneration. However, the molecular mechanism by which TDP-43 aggregates form and cause neurodegeneration remains poorly understood. Besides impairments in RNA and protein homeostasis, nucleocytoplasmic transport defects are emerging as converging disease mechanisms in ALS/FTD^8, 9^. Therapeutic interventions aiming to target the triad of TDP-43 control, i.e. autoregulation, nucleocytoplasmic transport and aggregation, is viewed as a promising strategy to prevent neurodegeneration^10^.

Cyclophilin A/peptidyl-prolyl *cis-trans* isomerase A (PPIA) is highly expressed in the CNS, where it localizes mainly in neurons^11^. Despite its abundance, its primary function in the CNS remains largely undefined. PPIA is involved in several cellular processes with double-edged functions. Inside the cells it is mainly beneficial. Besides promoting *de novo* protein folding^12^, it acts as a molecular chaperone and protects against oxidative stress and protein misfolding^13–15^. PPIA interacts with heterogeneous nuclear ribonucleoproteins (hnRNPs), regulates their nucleocytoplasmic transport and plays a role in the stability of the RNP complexes^13, 16^. PPIA interacts with TDP-43 and governs some of its functions^13^. In particular, it influences TDP-43 binding to its RNA targets affecting the expression of genes, such as *HDAC6*, *ATG7*, *VCP*, *FUS* and *GRN*, involved in the clearance of protein aggregates, and/or mutated in ALS and FTD^13^. Since PPIA Lys-acetylation favours the interaction with TDP-43, a post-translational regulation is hypothesized. Extracellularly, PPIA behaves as a pro-inflammatory cytokine^17^. Aberrantly secreted in response to stress, PPIA activates an EMMPRIN/CD147-NF-κB-MMP-9 pathway, promoting neuroinflammation and selective motor neuron death^18^.

Although PPIA has been studied in connection with several human diseases, some neurodegenerative, its role in pathogenesis has not been established^19^. We firstly reported that PPIA is altered in ALS^20–22^. In particular, we observed that PPIA accumulates in Triton-insoluble protein aggregates from the spinal cord in ALS mouse models and sporadic patients^20^. Next, we reported that low levels of soluble PPIA in PBMCs of ALS patients are associated with early onset of the disease^23^ and short disease duration^7^. In agreement with this, PPIA deficiency exacerbated aggregation and accelerated disease progression in a mutant SOD1 mouse model of ALS^13^. However, selective inhibition of extracellular PPIA protected motor neurons, reduced neuroinflammation and increased survival^18^.

In previous work we noted that PPIA knock-out (*PPIA-/-*) mice presented features of TDP-43 pathology but with no overt clinical phenotype up to four months of age^13^. Here we have characterized *PPIA-/-* mice neuropathologically and behaviourally throughout their entire lifespan and found that they develop a neurodegenerative disease with marked TDP-43 pathology. PPIA is also defective in several sporadic ALS and ALS-FTD patients, and a rare loss-of-function PPIA mutation was identified in an ALS patient, supporting its involvement in the etiopathogenesis.

## Materials and methods

### Animal model

Procedures involving animals and their care were conducted in conformity with the following laws, regulations, and policies governing the care and use of laboratory animals: Italian Governing Law (D.lgs 26/2014; Authorization 19/2008-A issued March 6, 2008 by Ministry of Health); Mario Negri Institutional Regulations and Policies providing internal authorization for persons conducting animal experiments (Quality Management System Certificate, UNIENISO9001:2008, Reg.No.6121); the National Institutes of Health’s Guide for the Care and Use of Laboratory Animals (2011 edition), and European Union directives and guidelines (EEC Council Directive, 2010/63/UE). The Mario Negri Institutional Animal Care and Use Committee and the Italian Ministry of Health (Direzione Generale della Sanità Animale e dei Farmaci Veterinari, Ufficio 6) prospectively reviewed and approved the animal research protocols of this study (prot. no. 14-02/C and 9F5F5.60) and ensured compliance with international and local animal welfare standards.

*PPIA-/-* mice were originally generated and characterized as described^24, 25^. We obtained *PPIA-/-* mice (strain 129S6/SvEvTac Ppiatm1Lubn/Ppiatm1Lbn; stock no. 005320) from The Jackson Laboratory; they were maintained on a 129S6/SvEvTac background. The *PPIA-/-* mice on 129S6/Sv genetic background and corresponding *PPIA+/+* littermates were used for micro-CT, MRI, immunohistochemistry, biochemistry, long-term potentiation, Rotarod, Grid test, Extension reflex, and will be indicated hereafter as *PPIA-/-* mice. *PPIA-/-* mice on C57BL/6J genetic background (C57_*PPIA-/-* mice), kindly provided by Dr. Bradford C. Berk (University of Rochester Medical Center, Rochester, New York, USA), and corresponding *PPIA+/+* littermates, were used for cognitive tests (open field, elevated-plus maze, three-chamber sociability, Morris water maze, novel object recognition). Animals were bred and maintained at the Istituto di Ricerche Farmacologiche Mario Negri IRCCS, Milano, Italy, as described in the Supplementary material.

### Micro-computed tomography (micro-CT) and MRI

Micro-CT and MRI analysis were done essentially as previously described^26, 27^. Further details are reported in the Supplementary material.

### Immunohistochemistry

Immunohistochemistry was done on coronal sections of brain for TDP-43, phosphorylated TDP-43 (pTDP-43), glial fibrillary acidic protein (GFAP), Iba1 and Nissl staining was done on brain and lumbar spinal cord sections, as described in the Supplementary material.

### Neurofilament light chain (NFL) measurements

Plasma samples were collected from *PPIA+/+* and *PPIA-/-* mice in K2-EDTA BD Microtainer blood collection tube and centrifuged at 5000 xg for 5 minutes to isolate plasma sample. The plasma NFL concentration was measured using the Simoa® NF-light™ Advantage (SR-X) Kit (#103400) on the Quanterix SR-X™ platform with reagents from a single lot, according to the protocol issued by the manufacturer (Quanterix Corp, Boston, MA, USA).

### Subcellular fractionation

Nuclear and cytoplasmic fractions were isolated from mouse cortex and cerebellum essentially as described^13, 18^. Further details are reported in the Supplementary material.

### Extraction of detergent-insoluble proteins

Mouse tissues were homogenized and the Triton-insoluble fraction (TIF) was obtained essentially as described in^13^. Quantification of the insoluble and soluble protein fractions was done as described in the Supplementary material.

### Immunoblotting

Western blot (WB) and dot blot were done as previously described^7, 20^. The antibodies used for immunoblot are reported in the Supplementary material.

### Real-time PCR

The total RNA from mouse cortex and human PBMC was extracted using the RNeasy® Mini Kit (Qiagen) and real-time PCR was done as described in the Supplementary material.

### Long-term potentiation (LTP) analysis

Coronal brain slices (350 µm) were cut and processed for LTP recordings as described in the Supplementary material.

### Behavioural analysis

In this study both male and female mice were tested, at 6 and 12 months of age. All behavioural tests were done at the same time of day, in the afternoon. Mice were allowed to habituate to the test room for at least 1h. Test environments were thoroughly cleaned between test sessions and males were tested before females. The open field, elevated-plus maze, three-chamber sociability, Morris water maze and novel object recognition test used Ethovision XT, 5.0 software (Noldus Information Technology, Wageningen, The Netherlands) to record the parameters. Further details are described in the Supplementary material.

### Muscle atrophy

Tibialis anterior muscles were freshly dissected, as previously described^28^, immediately frozen and weighted on an analytical balance with 0.1 mg readability and ± 0.2 mg linearity (Crystal series, Gibertini). To evaluate muscle atrophy, the weight of the muscle was normalized to mouse body weight.

### Subjects in the study

Informed written consent was obtained from all subjects involved and the study was approved by the ethics committee of Azienda Ospedaliero Universitaria Città della Salute e della Scienza, Turin and ICS Maugeri IRCCS, Milano. The diagnosis of ALS was based on a detailed medical history and physical examination, and confirmed by electrophysiological evaluation. Inclusion criteria for ALS patients were: i) 18-85 years old; ii) diagnosis of definite, probable or laboratory-supported probable ALS, according to revised El Escorial criteria^29^. Exclusion criteria were: i) diabetes or severe inflammatory conditions; ii) active malignancy; iii) pregnancy or breast-feeding. ALS patients underwent a battery of neuropsychological tests and were classified according to the consensus criteria for the diagnosis of frontotemporal cognitive and behavioural syndromes in ALS^30–32^. We analysed clinical samples from three independent cohorts of ALS patients and controls (Cohort #1-3), totally 151 ALS patients and 128 healthy subjects. The characteristics of the patients and controls are described in the Supplementary Table 1.

### Isolation of PBMCs and plasma

Blood was drawn by standard venipuncture into Vacutainer® Plus Plastic K2EDTA Tubes (Becton, Dickinson and Company) and kept at 4°C until shipment to the Istituto di Ricerche Farmacologiche Mario Negri IRCCS. PBMCs were isolated from ALS patients and healthy individuals, as previously described^7^. Further details are described in the Supplementary material.

### Mutation screening

We examined a cohort of 959 ALS patients from Northern Italy and 677 healthy controls matched by age, sex and geographical origin. Informed written consent was obtained for all subjects and the study was approved by the ethics committees involved. Whole-genome sequencing was done at The American Genome Center (Uniformed Services University, Walter Reed National Military Medical Center campus, Bethesda, MD, USA) as described in the Supplementary material.

### Molecular dynamics (MD) simulations

The structure of WT PPIA was retrieved from PDB 1CWA. The mutant form was generated by swapping the lysine 76 with glutamate using UCSF Chimera^33^. MD simulations were carried out as described in the Supplementary material.

### Comparative analysis with retrospective cohort

To compare protein levels of the PPIA K76E patient with the retrospective cohort analysed in Luotti et al. 2020^7^, PBMCs were isolated and proteins extracted exactly as previously reported^7^. Slot blot analysis was done with the same internal standard (IS) as the previous work, which is a pool of all samples in the analysis. Immunoreactivity was normalized to Red Ponceau staining (Fluka) and to the immunosignal of the IS.

### AlphaLISA assay of MMP-9

The level of MMP-9 in plasma was measured with an AlphaLISA kit for the human protein (#AL3138, PerkinElmer). AlphaLISA signals were measured using an Ensight Multimode Plate Reader (PerkinElmer).

### Cell and molecular biology procedures

Mutant PPIA cloning, cell culture, transfection and treatments were done as described in the Supplementary material.

### Immunoprecipitation

Magnetic beads coupled with sheep polyclonal antibodies anti-rabbit IgG (Dynabeads, Invitrogen) were used for co-immunoprecipitation studies. Cells were lysed in 50 mM Tris-HCl, pH 7.2, 2% CHAPS, protease inhibitor cocktail (Roche) and quantified with the BCA protein assay (Pierce). Proteins (500 µg) were diluted to 0.5 µg/µL with lysis buffer. Magnetic beads with coupled sheep antibodies anti-rabbit IgG (Dynabeads® M280; Invitrogen) were washed with 0.5% bovine serum albumin immonuglobulin-free (BSA Ig-free) in PBS to remove preservatives. Three μg of rabbit polyclonal anti-human TDP-43 primary antibody (Proteintech; RRID: AB_2200505) or rabbit polyclonal anti-human Ran primary antibody (Abcam) was incubated with 20 μL of anti-rabbit IgG-conjugated Dynabeads for 2h at 4°C in 0.1% BSA/PBS. Immunoprecipitation and analysis of the proteins are described in the Supplementary material.

### Statistical analysis

Prism 7.0 (GraphPad Software Inc., San Diego, CA) was used. For each variable, the differences between experimental groups were analysed by Student’s t test, or one-way ANOVA followed by post-hoc tests. Pearson’s correlation coefficient (r) was calculated to measure the linear strength of the association between PPIA and TDP-43 total protein levels. Two-way ANOVA followed by the Bonferroni post-hoc test was used to analyse Rotarod performance and compare the body weight of PPIA -/- mice with controls. Survival curves were analysed using a log-rank Mantel Cox test. P values below 0.05 were considered significant.

### Data availability

The whole genome sequence data is publicly available on dbGaP at phs001963. The other data that support the findings of this study are available within the manuscript and supplementary material. Source data are available from the corresponding author upon reasonable request.

## Results

### PPIA is defective in sporadic ALS patients

In PBMCs of ALS patients we detected a low level of Lys-acetylated PPIA^13^. We also observed, an impaired soluble/insoluble partitioning of PPIA and TDP-43 in PBMCs of a relatively large cohort of sporadic ALS and ALS-FTD patients in comparison with healthy controls, with significant accumulation of PPIA and TDP-43 in the insoluble fraction and substantially less PPIA in the soluble fraction^7^. Here we measured the total PPIA and TDP-43 protein levels in the retrospective cohort (Cohort #1, Supplementary Table 1). PPIA was significantly lower than controls, while TDP-43 did not differ (Fig. 1A-B). We checked whether this reflected downregulation of the PPIA gene. There was in fact significant downregulation of PPIA at the mRNA level too, in an independent cohort of sporadic ALS patients compared to healthy controls (Cohort #2, Supplementary Table 1), and only a tendency to lower TDP-43 mRNA (Fig.1C-D). Finally, we detected a significant, although weak, positive correlation (r=0.3, p=0.01) between PPIA and TDP-43 total protein levels.

**Figure 1.**
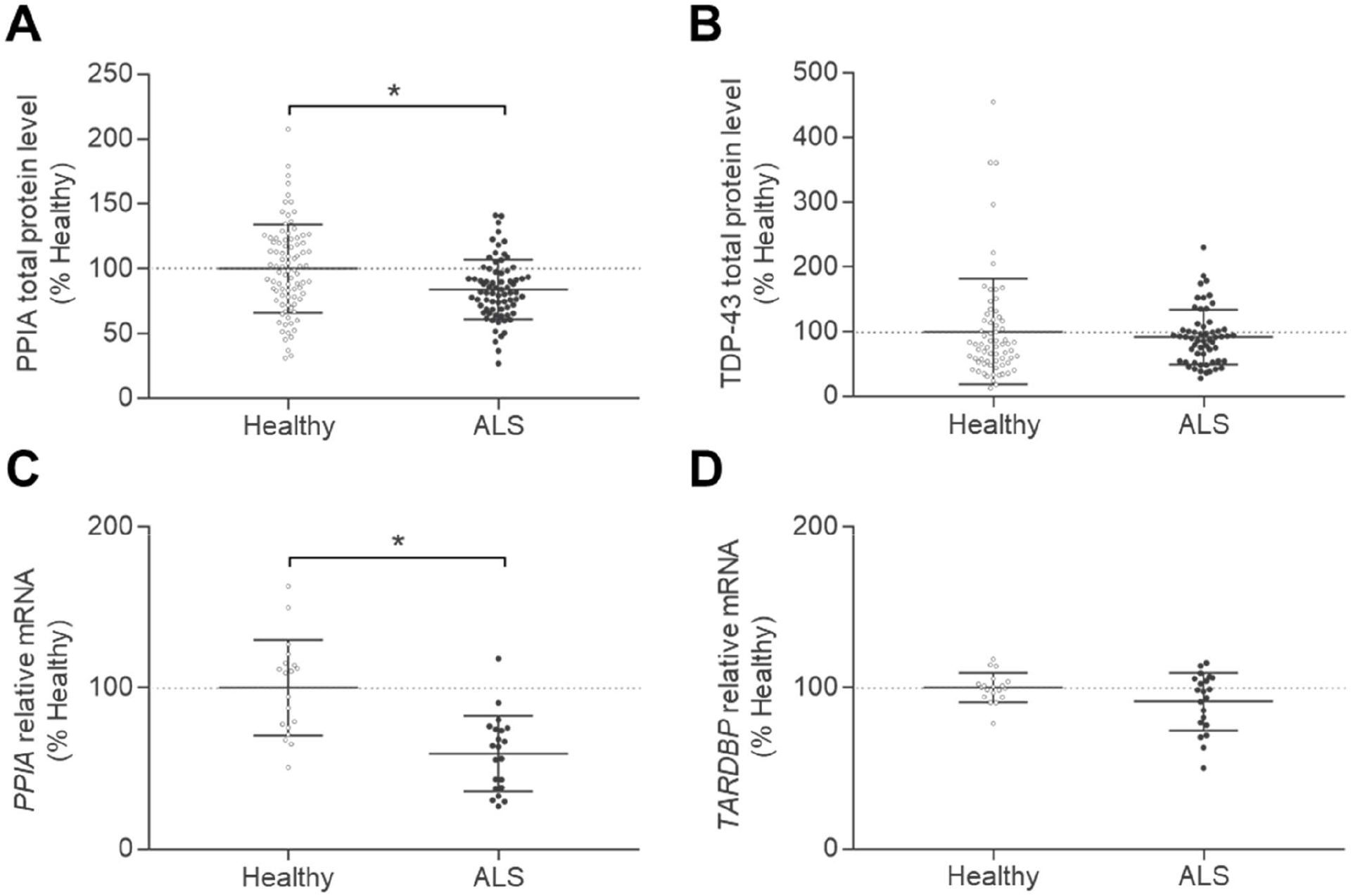
PPIA is downregulated in sporadic ALS patients. (**A,B**) Total PPIA (**A**) and TDP-43 **(B)** protein levels were analysed in PBMCs from ALS patients and healthy subjects of a retrospective cohort^7^ (Cohort #1, Supplementary Table1). Scatter dot plots (mean ± SD; PPIA: n=89 healthy controls, n=74 ALS patients; TDP-43: n=69 healthy controls, n=65 ALS patients) are percentages of healthy controls and the dotted lines indicate the median of healthy controls. *p <0.05 versus healthy controls, Student’s t-test. **(C,D)** Real-time PCR for PPIA **(C)** and *TARDBP* **(D)** mRNA transcripts in PBMCs of ALS patients and healthy subjects (Cohort #2, Supplementary Table1). Data (mean ± SD; n=19 healthy subjects; n=21 ALS patients) are normalized to β-actin and expressed as percentages of the healthy control relative mRNA. Dotted lines indicate the median of healthy controls. *p <0.05 versus healthy controls, Student’s t-test

In conclusion, defective PPIA is a common feature of ALS and ALS-FTD patients and could trigger TDP-43 pathology.

### *PPIA-/-* mice display neuropathological alterations, worsening with age

To explore PPIA’s impact on TDP-43 biology we did deep characterization of *PPIA-/-* mice throughout their lifespan. In our previous studies *PPIA-/-* mice had a higher burden of Triton-insoluble phosphorylated TDP-43 (insoluble pTDP-43) in the spinal cord and brain cortex already at four months of age^13^. We detected no motor neuron loss^18^ and no motor phenotype (Supplementary Fig. 1A-B), but there was a tendency to kyphosis (Supplementary Fig. 1C).

To see whether these were prodromal signs of neurodegeneration, we did a neuropathological analysis of older *PPIA-/-* mice. Quantitative MRI was done longitudinally on *PPIA-/-* and *PPIA+/+* mice at 6 and 12 months. *PPIA-/-* mice had a smaller total brain volume than controls, with the most marked difference at 12 months (−13%) (Supplementary Fig. 1D).

Next, we verified the effect of PPIA depletion on hippocampus, cortex and cerebellum, adjusting for total brain volume (Fig. 2 and Supplementary Fig. 1). *PPIA-/-* mice had a significantly smaller hippocampus than *PPIA+/+* mice at 6 and 12 months (Fig. 2A). There was also a reduction of the relative volume with age, independently from the genotype, slightly more marked in *PPIA-/-* mice (−23% versus -21%). The cortex volume of *PPIA-/-* mice was also smaller than in *PPIA+/+* mice, but less so: 3% and 10% smaller respectively at 6 and 12 months of age, but the difference was significant only at 12 months (Fig. 2B). However, *PPIA-/-* mice had significant cortex atrophy at 12 compared to 6 months (−16%); this was less marked in control mice (−10%). In contrast, cerebellum relative volume did not significantly differ in *PPIA-/-* mice and controls and did not change with age (Supplementary Fig. 1E).

**Figure 2.**
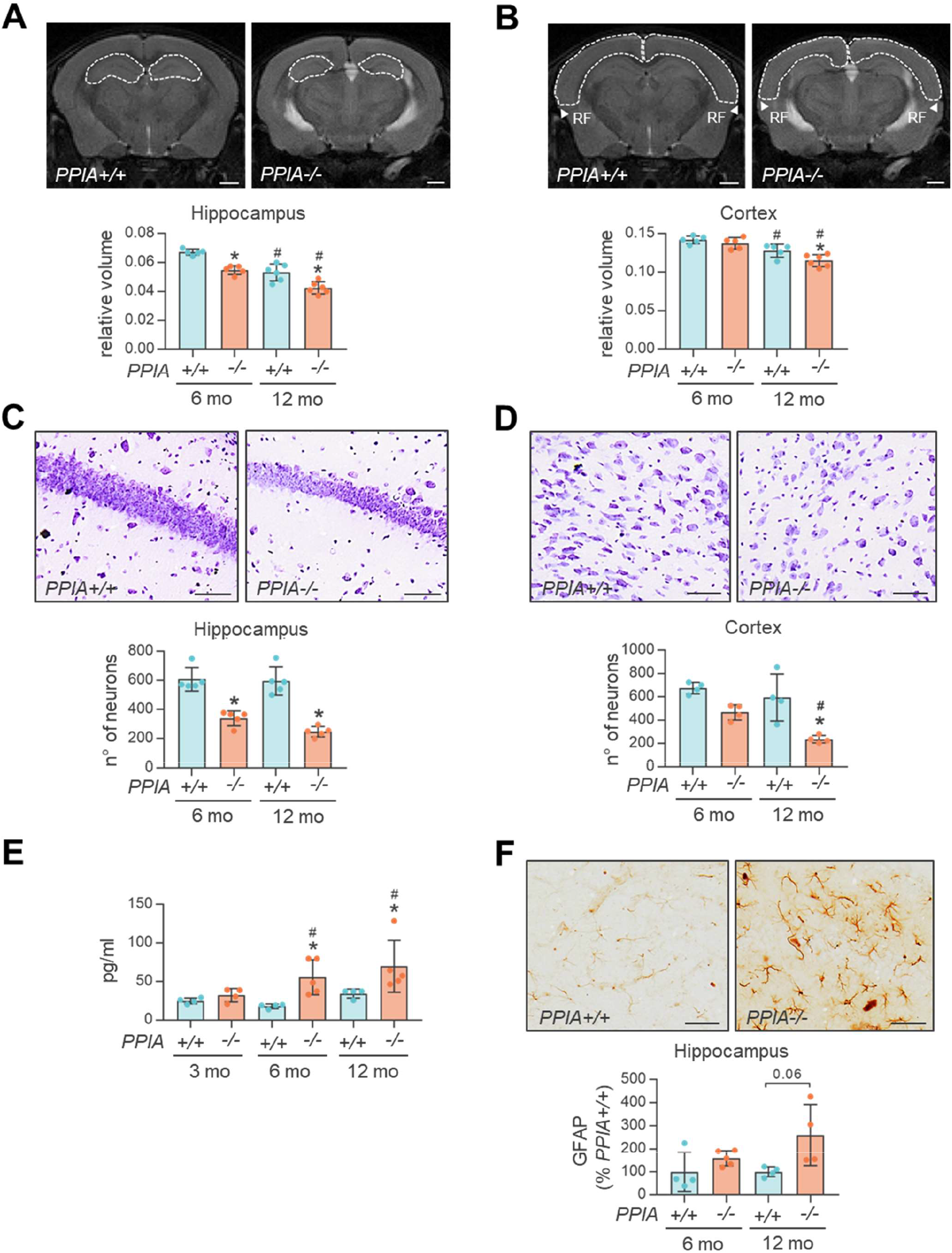
*PPIA-/-* mice show progressive neuropathological alterations. **(A,B)** The volume of hippocampus **(A)** and cortex **(B)** was measured using quantitative MRI in *PPIA+/+* and -/- mice, at 6 and 12 months (mo) of age. Representative MRI images of *PPIA+/+* and -/- brain regions at 12 months are shown. The white dashed lines indicate the ROI considered for MRI quantification. Scale bar 1 mm. The volume of hippocampus and cortex were adjusted for total brain volume and data are expressed as relative volume: mean ± SD (n=5 or 6 in each experimental group). **(C,D)** Depletion of PPIA induces progressive neuron loss. Neurons in CA1 region **(C)** and cortex **(D)** were counted in *PPIA+/+* and -/- mice at 6 and 12 months. Representative Nissl-stained brain section are shown. Scale bar 50 µm. Mean ± SD (n=5, CA1 region; n=4, cortex). **(E)** NFL plasma levels were analysed in *PPIA+/+* and *PPIA-/-* mice at 3, 6 and 12 months. Data are mean ± SD (n=4 or 5 in each experimental group). *p >0.05 versus *PPIA+/+* mice and #p <0.05 versus the 3 months, by Student’s t-test. **(F)** GFAP-immunostaining in hippocampus was quantified in *PPIA+/+* and -/- mice at 6 and 12 months. Representative GFAP-stained brain sections are shown. Scale bar 50 µm. Data (mean ± SD; n=4 or 5 in each experimental group) are percentages of the *PPIA+/+*. **(A-D, F)** *p <0.05 versus the *PPIA+/+* mice and ^#^p <0.05 versus the 6 months, by one-way ANOVA, Tukey’s post hoc test.

To investigate whether brain atrophy reflected neuronal loss we did a histological analysis on the hippocampus and cortex (Fig. 2C-D). There were substantially fewer Nissl-positive neurons in the hippocampus CA1 region of *PPIA-/-* mice than in controls at 6 and 12 months, and this tended to be more pronounced at 12 months (Fig. 2C). In the cortex *PPIA-/-* mice had fewer Nissl-positive neurons than controls at 6 and 12 months, but this was significant only at 12 months (Fig. 2D). They also had significantly greater neuron loss at 12 months than at 6 months, similarly to the MRI picture. The progressive neurodegenerative process in *PPIA-/-* mice was also confirmed by the plasma NFL level, a marker of neuro-axonal damage, that increased substantially with age compared to controls from 6 months (Fig. 2E).

We next examined the glial response in the hippocampus and cortex of *PPIA-/-* mice (Fig. 2F, Supplementary Fig. 1F-G). In the hippocampus, PPIA deficiency tended to increase astroglial activation at 12 months (Fig. 2F), but had no effect on microglia (Supplementary Fig. 1F). In the cortex, PPIA deficiency did not affect either astroglia or microglia (Supplementary Fig. 1G), confirming PPIA role in microglial activation and the neuroinflammatory response^18^.

To explore whether the neuropathological alterations were associated with TDP-43 pathology, we did biochemical and immunohistochemistry analyses for TDP-43 and pTDP-43 in brains of *PPIA-/-* mice and controls at 6 and 12 months of age (Fig. 3, Supplementary Fig. 2). In the cortex of *PPIA-/-* mice there was a significant increase in pTDP-43 in the cytoplasm and a concomitant decrease of TDP-43 in the nucleus compared to controls already at 6 months (Fig. 3A), while we saw no change in cerebellum (Supplementary Fig. 2A). We analysed the Triton-insoluble protein fraction from brain cortex of *PPIA-/-* and *PPIA+/+* mice and found that in *PPIA-/-* mice there was significantly larger amount of insoluble proteins, TDP-43 and pTDP-43 (Supplementary Fig. 2B-D). We also analysed other proteins associated with ALS/FTD that interact with TDP-43 in RNP complexes and/or stress granules (Supplementary Fig. 2E-F). In the absence of PPIA, there was a higher level of insoluble hnRNPA2/B1 and TIA1, confirming the negative effect of deficient PPIA folding/refolding activity in the brain^13^.

**Figure 3.**
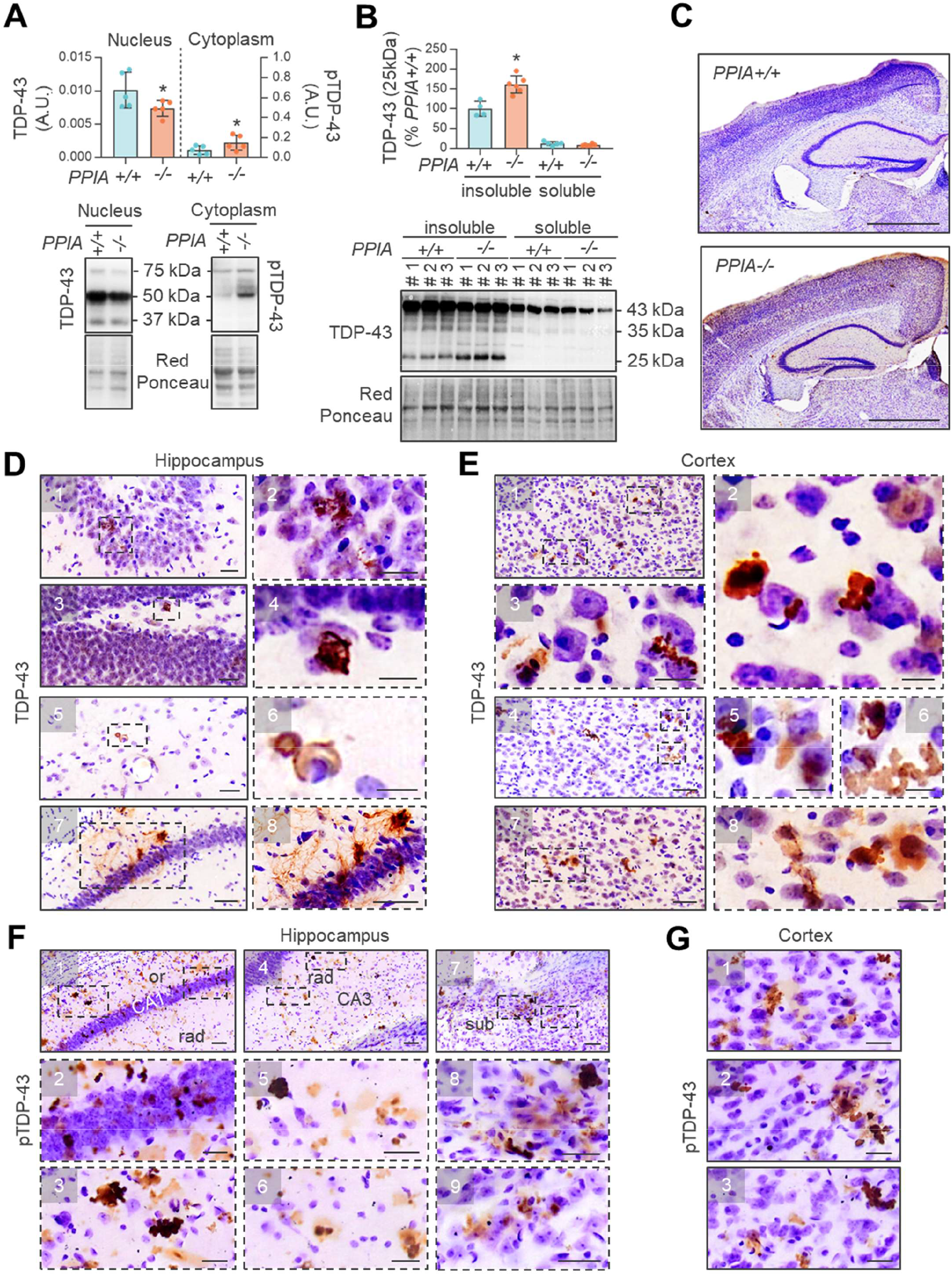
TDP-43 pathology in hippocampus and cortex of *PPIA-/-* mice. TDP-43 pathology was analysed by biochemical **(A,B)** and histological **(C-G)** approaches, at 6 and 12 months. **(A)** Equal amounts of nuclear and cytoplasmic fractions from cortex of *PPIA+/+* and -/- mice were analysed by WB for TDP-43 (left) and pTDP-43 (right). Representative WB are shown. Data (mean ± SD; n=5 in each experimental group) indicates the immunoreactivity normalized to total protein loading, in arbitrary units (A.U.). *p < 0.05 versus *PPIA+/+* mice Student’s t-test. **(B)** Representative WB of TDP-43 in soluble and insoluble fractions from cortex of *PPIA+/+* and -/- mice and quantification of the 25 kDa TDP-43 fragment are shown. Data (mean ± SD, n=4-6 in each experimental group) are percentages of immunoreactivity in *PPIA+/+* insoluble fraction. *p < 0.05 versus the *PPIA+/+* mice by one-way ANOVA, Tukey’s post hoc test. **(C)** Diffuse TDP-43 immunostaining was observed in brains of *PPIA-/-* mice at 12 months compared to *PPIA+/+*. Scale bar 1 mm. **(D)** In hippocampus of *PPIA-/-* mice, there were TDP-43 inclusions in the pyramidal layer of CA3 and CA1 (1-2), in the granule cell layer of dentate gyrus (3-4), in stratum radiatum of CA1 (5-6) and in stratum oriens of CA3 and CA1 (7-8). Panels 2,4,6,8 are magnified images of the dashed area in panels 1,3,5,7. Scale bar 50 µm in panels 7,8; scale bar 25 µm in panels 1,3,5; scale bar 20 µm in panels 2; scale bar 10 µm in panels 4,6. **(E)** *PPIA-/-* mice had TDP-43 inclusions in the somatosensory (1-6) and temporal cortex (7-8). Panels 2,3,5,6,8 are magnified images of the dashed area in panels 1,4,7. Scale bar 50 µm in panels 1,4,7; scale bar 20 µm in panels 3,8; scale bar 10 µm in panels 2,5,6. **(F,G)** pTDP-43 immunostaining was analysed in hippocampus **(F)** and cortex **(G)** of *PPIA-/-* mice and control *PPIA+/+* mice (Supplementary Fig. 2G-H) at 12 months. **(F)** pTDP-43 inclusions are widespread in the CA1 pyramidal layer (1-2), stratum oriens (or) of CA1 (3), stratum radiatum (rad) of CA3 (4-6) and subiculum (sub) (7-9). Panels 2,3,5,6,8,9 are magnified images of the dashed area in panels 1,4,7. Scale bar 50 µm in panels 1,4,7; scale bar 25 µm in panels 5,8,9; scale bar 20 µm in panels 2,3,6. **(G)** pTDP-43 staining was observed in the somatosensory (1) and auditory-temporal (2-3) cortex. Scale bar 20 µm.

Finally, we examined TDP-43 fragmentation in soluble and insoluble brain cortex fractions (Fig. 3B). Characteristic C-terminal 35- and 25-kDa TDP-43 fragments were abundant, mainly in the insoluble fraction. While TDP-43 bands at 43 and 35 kDa respectively slightly decrease (−6%) and increase (+12%) in *PPIA-/-* mice, but not significantly (data not shown), the 25-kDa TDP-43 fragment accumulated substantially (+61%). As often happens in ALS/FTD patients^34, 35^, the burden of TDP-43 C-terminal fragments is greater than that of TDP-43 full-length and the 25 kDa accumulates more than the 35 kDa fragment, probably because is less readily degraded^36^.

To better explore TDP-43 pathology and C-terminal TDP-43 fragments we did an histopathological analysis with an antibody that targets the C-terminus of the protein^37^. At 12 months of age, there was a widespread TDP-43 signal in different brain regions of *PPIA-/-* mice, with higher intensity in hippocampus and cortex, that accumulates in aggregated structures (Fig. 3C-E). We confirmed the presence of TDP-43 inclusions in *PPIA-/-* brain regions using a pTDP-43 antibody that targets the C- terminus, does not react with physiological nuclear TDP-43 but identifies only pathological brain lesions (Fig. 3F-G and Supplementary Fig. 2G-H).

We concluded that PPIA deficiency induces a clear-cut neuropathological phenotype in the mouse brain, with marked TDP-43 pathology and other alterations related to protein and RNA homeostasis, getting worsen with age.

### PPIA deficiency affects proteins involved in nucleocytoplasmic transport, TDP-43 autoregulation and synaptic function

PPIA is a foldase and a molecular chaperone potentially for a wide range of substrates and interacts with GTP-binding nuclear protein Ran (Ran)^13^. TDP-43 requires functional Ran for import into the nucleus and regulates its expression^38^. In the cortex of *PPIA-/-* mice, Ran total protein level did not differ from controls (data not shown). However, Ran accumulated substantially (+69%) in the detergent-insoluble fraction and decreased in the soluble fraction (−38%) (Fig. 4A), already at 6 months of age. These findings may indicate that depletion of the folding/refolding activity of PPIA is at the basis of the reduced solubility of Ran and contribute to defective nucleocytoplasmic transport.

**Figure 4.**
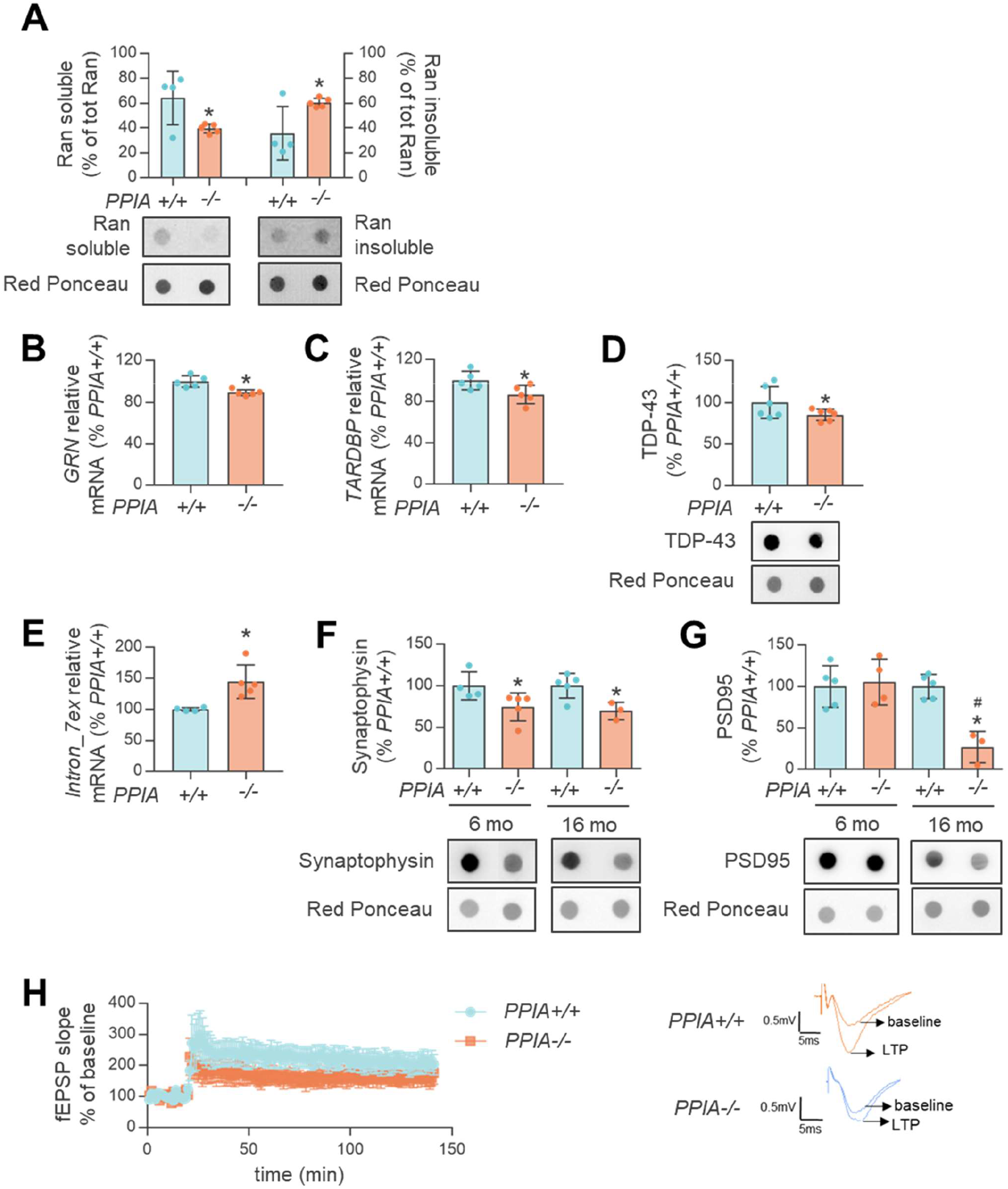
PPIA deficiency affects proteins involved in nucleocytoplasmic transport and synaptic function. **(A)** Dot blot analysis for Ran-soluble and -insoluble protein in cortex of *PPIA+/+* and *PPIA-/-* mice at 6 months of age. Data (mean ± SD; n=4 or 5 in each experimental group) are normalized to protein loading and expressed as percentages of total protein, of Ran in soluble and insoluble fractions. Representative dot blots are shown. (B-D) Real-time PCR for GRN **(B)** and TARDBP **(C)** mRNA transcripts and dot blot analysis for TDP-43 total protein **(D)**, in cortex of *PPIA+/+* and *PPIA-/-* mice at 6 months. **(B,C)** Data (mean ± SD; n=5) are normalized to β-actin and expressed as percentages of *PPIA+/+* relative mRNA. **(D)** Data (mean ± SD, n=6) are normalized to protein loading and are percentages of immunoreactivity in *PPIA+/+* mice. Representative dot blots are shown. **(E)** Real-time PCR for TARDBP intron 7 exclusion in cortex of *PPIA+/+* and *PPIA-/-* mice at 6 months. Data (mean ± SD; n=4 or 5 in each experimental group) are normalized to β-actin and expressed as percentages of *PPIA+/+* relative mRNA. (A-E) *p < 0.05 versus *PPIA+/+* mice, Student’s t-test. **(F,G)** Dot blot analysis of synaptophysin **(F)** and PSD95 **(G)** in cortex of *PPIA+/+* and -/- mice at 6 and 16 months. Immunoreactivity was normalized to protein loading. Representative dot blots are shown. Data (mean ± SD, n=3 or 5 in each experimental group) are percentages of immunoreactivity in *PPIA+/+* mice. *p <0.05 versus *PPIA+/+* mice and #p <0.05 versus 6 months, by one-way ANOVA, uncorrected Fisher’s LSD post hoc test **(F)** and Tukey’s post hoc test **(G)**. **(H)** CA1 long-term potentiation (LTP) induced with theta burst stimulation (TBS) was lower in *PPIA-/-* mice (white rectangle) than *PPIA+/+* mice (black circles) at 16 months. Data were analysed with two-way ANOVA for repeated measures, p < 0.05 (n=6). Representative traces recorded in slices from *PPIA+/+* and *PPIA-/-* mice, before and after TBS, are shown.

PPIA deficiency affects the expression of a number of TDP-43 RNA targets, including GRN^13^. GRN mutations in patients result in haplo-insufficiency and GRN knock-out mice present an FTD-like phenotype associated with synaptic dysfunction^39^. We verified whether knocking out PPIA influenced GRN expression in the brain cortex of the mice. GRN mRNA levels were slightly but significantly lower in *PPIA-/-* mice than controls (Fig. 4B). Surprisingly, TDP-43 at mRNA and protein levels was also lower in *PPIA-/-* mice than controls (Fig. 4C-D). TDP-43 adjusts its own expression by binding sequences within the 3′ UTR of its own transcript, promoting processing of alternative introns (e.g. intron 7) and mRNA destabilization^2, 3^. We found a 45% increase in the exclusion of *TARDBP* intron 7 in *PPIA-/-* compared to *PPIA+/+* mice (Fig. 4E). This suggests that PPIA deficiency may also interfere with the mechanism of TDP-43 autoregulation leading to TDP-43 downregulation.

TDP-43 depletion in brain downregulates genes involved in synaptic function^3^. To assess whether GRN and *TARDBP* downregulation was associated with synaptic dysfunction, we first measured the levels of synaptophysin and PSD95, markers of pre- and post-synaptic structures, in the cortex of *PPIA-/-* mice and controls at 6 and 16 months of age (Fig. 4F-G). Both synaptophysin and PSD95 were lower in *PPIA-/-* than *PPIA+/+* mice. Next, we made extracellular field recordings of excitatory postsynaptic potentials in the CA1 hippocampal region of *PPIA-/-* mice at 16 months of age compared to controls. Long-term potentiation induced with theta burst stimulation had a smaller amplitude in *PPIA-/-* mice (Fig. 4H), suggesting impairment of synaptic plasticity in the hippocampus.

### *PPIA-/-* mice develop cognitive, behavioural impairments and late-onset motor neuron disease

We checked whether PPIA deficiency, besides compromising neuronal functions, promotes cognitive, behavioural and motor impairments. Since mice on a 129S6/Sv genetic background do not perform well in cognitive tasks^40^, C57_*PPIA-/-* mice were used for these experiments. We first confirmed that also C57_*PPIA-/-* mice display TDP-43 pathology (Supplementary Fig. 3). Next, we found that C57_*PPIA-/-* mice showed no exploratory abnormalities, as they spent the same time as C57_*PPIA+/+* mice in the inner zone in the open field test, at both 6 and 12 months of age (Fig. 5A). In the elevated-plus maze paradigm C57_*PPIA-/-* mice showed less anxiety and more disinhibition as they spent twice the time in the open arms compared with controls (Fig. 5B). This behaviour is lost with time.

**Figure 5.**
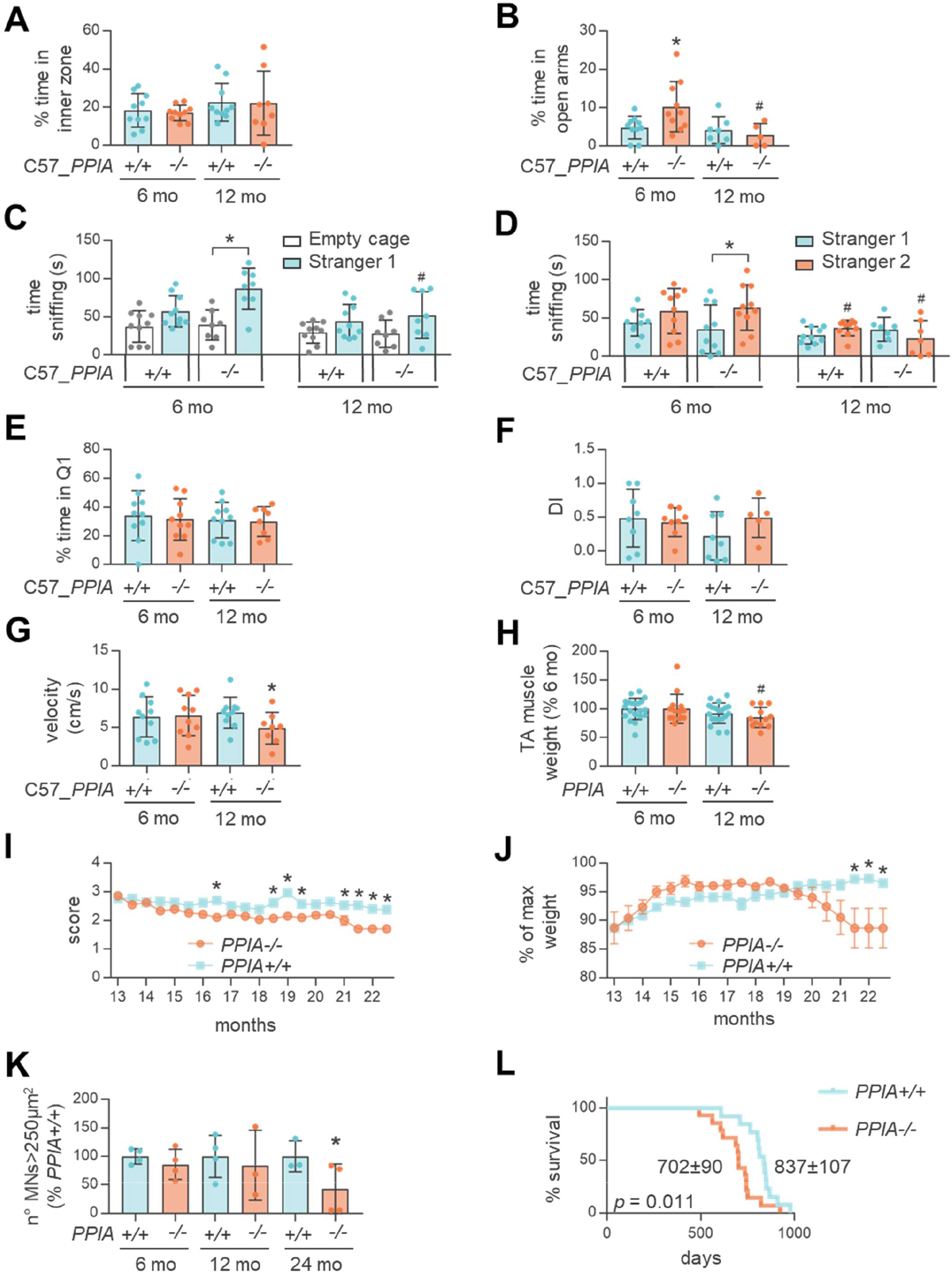
*PPIA-/-* mice present cognitive, behavioural and motor function deficits. **(A)** Open Field: C57_*PPIA+/+* and -/- mice spent similar time in the inner zone both at 6 (n=10) and 12 months of age (n=10 C57_*PPIA+/+*, n=8 C57_*PPIA-/-*). **(B)** Elevated plus maze: C57_*PPIA-/-* mice spent more time than controls in open arms (n=10 C57_*PPIA+/+* and -/- at 6 months, n=7 C57_*PPIA+/+* and n=5 C57_*PPIA-/-* at 12 months) at 6, but not at 12 months of age. **(C-D)** Three-chamber sociability test. In the sociability trial **(C)**, C57_*PPIA-/-* mice spent more time than controls sniffing Stranger 1 than Empty cage at 6, but not 12 months of age (n=10 C57_*PPIA+/+*, n=8 C57_*PPIA-/-*). In the social memory trial **(D)**, C57_*PPIA-/-* mice spent more time sniffing Stranger 2 than Stranger 1, compared to controls, at 6 (n=10), but not at 12 months (n=10 C57_*PPIA+/+*, n=7 C57_*PPIA-/-*). **(E)** Morris water maze: C57_*PPIA-/-* and +/+ mice spent similar time in the target quadrant (Q1) both at 6 (n=10) and 12 (n=10 C57_*PPIA+/+*, n=8 C57_*PPIA-/-*) months. **(F)** Novel object recognition test: C57_*PPIA+/+* and -/- mice had similar discrimination indexes (DI) at 6 (n=8) and 12 (n=8 C57_*PPIA+/+*, n=5 C57_*PPIA-/-*) months of age. **(A-F)** Mean ± SD. *p < 0.05 versus the *PPIA+/+* mice and ^#^p < 0.05 versus the 6-month-old control by one-way ANOVA, Tukey’s post hoc test **(A-C, E, F)** and uncorrected Fisher’s LSD post hoc test **(D)**. **(G)** Velocity measurement: C57_*PPIA+/+* and -/- mice had the same velocity in the open field test at 6 months (n=10) but at 12 months (n=10 C57_*PPIA+/+*, n=8 C57_*PPIA-/-*) C57_*PPIA-/-* had slower velocity than controls. Data are mean ± SD and are expressed in cm/s. *p <0.05 versus C57_*PPIA+/+* mice, Student’s t-test. **(H)** *PPIA-/-* mice show mild atrophy of tibialis anterior (TA) muscle at 12 months (n=20 *PPIA+/+*, n=14 *PPIA-/-* at 6 months, n=12 *PPIA-/-* at 12 months). Data (mean ± SD) are muscle weights as percentage of 6-month-old control. #p <0.05 versus 6-month-old control by Student’s t-test. **(I)** Extension reflex: *PPIA-/-* mice had less extension of hindlimbs than controls (n=13 *PPIA+/+*, n=14 *PPIA-/-*). Mean ± SEM. *p <0.05 by two-way ANOVA, Bonferroni’s post hoc test. **(J)** *PPIA-/-* mice had greater weight loss than controls from 21 months (n=13 *PPIA+/+*, n=14 *PPIA-/-*). Mean ± SEM. *p <0.05 by two-way ANOVA, Bonferroni’s post hoc test. **(K)** Quantification of Nissl-stained motor neurons (MNs > 250 μm^2^) in lumbar spinal cord hemisections from *PPIA+/+* and -/- mice at 6, 12 and 24 months. Data are mean ± SD (n=3 or 4 in each experimental group) and are percentages of *PPIA+/+* mice. *p <0.05 versus *PPIA+/+* mice, Student’s t-test. **(l)** Kaplan-Meier curve for survival of *PPIA+/+* (n=13) and *PPIA-/-* mice (n=14). Log-rank Mantel-Cox test for comparing *PPIA+/+* and -/- mice.

We used the three-chamber test to examine sociability and social memory in C57_*PPIA-/-* mice. C57_*PPIA-/-* mice at 6 months spent twice the time of controls sniffing the stranger than the empty cage (Fig. 5C), suggesting a more social attitude. However, at 12 months C57_*PPIA-/-* mice behaved differently, spending the same time with the stranger and the empty cage and significantly less time sniffing the stranger compared with mice at 6 months.

Combining the results of the elevated-plus maze and the three-chamber tests suggests that the greater social attitude of C57_*PPIA-/-* mice may be the consequence of greater disinhibition. During the social recognition memory task, which requires normal hippocampal function, C57_*PPIA-/-* mice at 6 months distinguished the stranger mouse (Stranger 2) from the familiar one (Stranger 1) (Fig. 5D), suggesting a normal social memory. However, at 12 months they no longer did it, suggesting social impairment and hippocampal dysfunction.

To investigate the role of impaired hippocampus in C57_*PPIA-/-* mice further, we used the Morris water maze and novel object recognition tests. C57_*PPIA-/-* and control mice showed no differences (Fig. 5E-F), suggesting no memory impairment. However, these tests were developed to study Alzheimer’s disease-related memory defects in mice and specific tests for FTD are not available^41^. Furthermore, also FTD patients show mild memory impairments despite the hippocampal degeneration^42^.

At 12 months C57_*PPIA-/-* mice had lower locomotor speed (Fig. 5G). In *PPIA-/-* mice, there was no evident motor impairment, as detected by rotarod and grid test throughout the lifespan (Supplementary Fig. 4A-B). However, at 12 months *PPIA-/-* mice had mild muscle atrophy (Fig. 5H), from 16 months impaired extension reflex (Fig. 5I), and from 21 months loss of body weight (Fig. 5J). These results correlated with gradual loss of motor neurons in the lumbar spinal tract of *PPIA-/-* mice, that became significant in the final stage of the disease (Fig. 5K). Finally, there was more than four months shorter survival of *PPIA-/-* mice than *PPIA+/+* mice (702 ± 90 days versus 837 ± 107 days) (Fig. 5L).

In summary, mice knock-out for PPIA present cognitive, behavioural and motor impairments reminiscent of ALS-FTD in patients.

### Identification of a PPIA loss-of-function mutation in a sporadic ALS patient

We identified an ALS patient with an heterozygous PPIA missense mutation (NM_021130: c.226A>G.; p.K76E) in an evolutionarily conserved region of the protein (Fig. 6A). To our knowledge, the PPIA:p.K76E is a novel variant absent in the large population dataset gnomAD v2.1.1, in our internal control cohort of 677 healthy subjects, matched by age and ancestry, and in the large ALS/FTD dataset ProjectMine (http://databrowser.projectmine.com/). The patient had difficulty walking at age 56 and slowly progressed to weakness and spasticity of the lower limbs. Neurological examination revealed increased deep tendon reflex in all extremities and pathological reflexes, including bilateral Hoffman sign and Babinski sign. No additional neurologic or systemic abnormalities were detected. The electromyography showed signs of lower motor neuron involvement in the upper and lower limbs. Genetic screening for mutations in the most common ALS genes and in a large panel of hereditary spastic paraplegia genes was negative. Appropriate investigations excluded other diseases, and motor impairment progressed over time. He was diagnosed with ALS at age 59. Cognitive testing was normal. No weakness in the upper limbs or bulbar and respiratory symptoms was reported after five years. The patient reported no family history of neurodegenerative disease, psychiatric disease or walking impairment.

**Figure 6.**
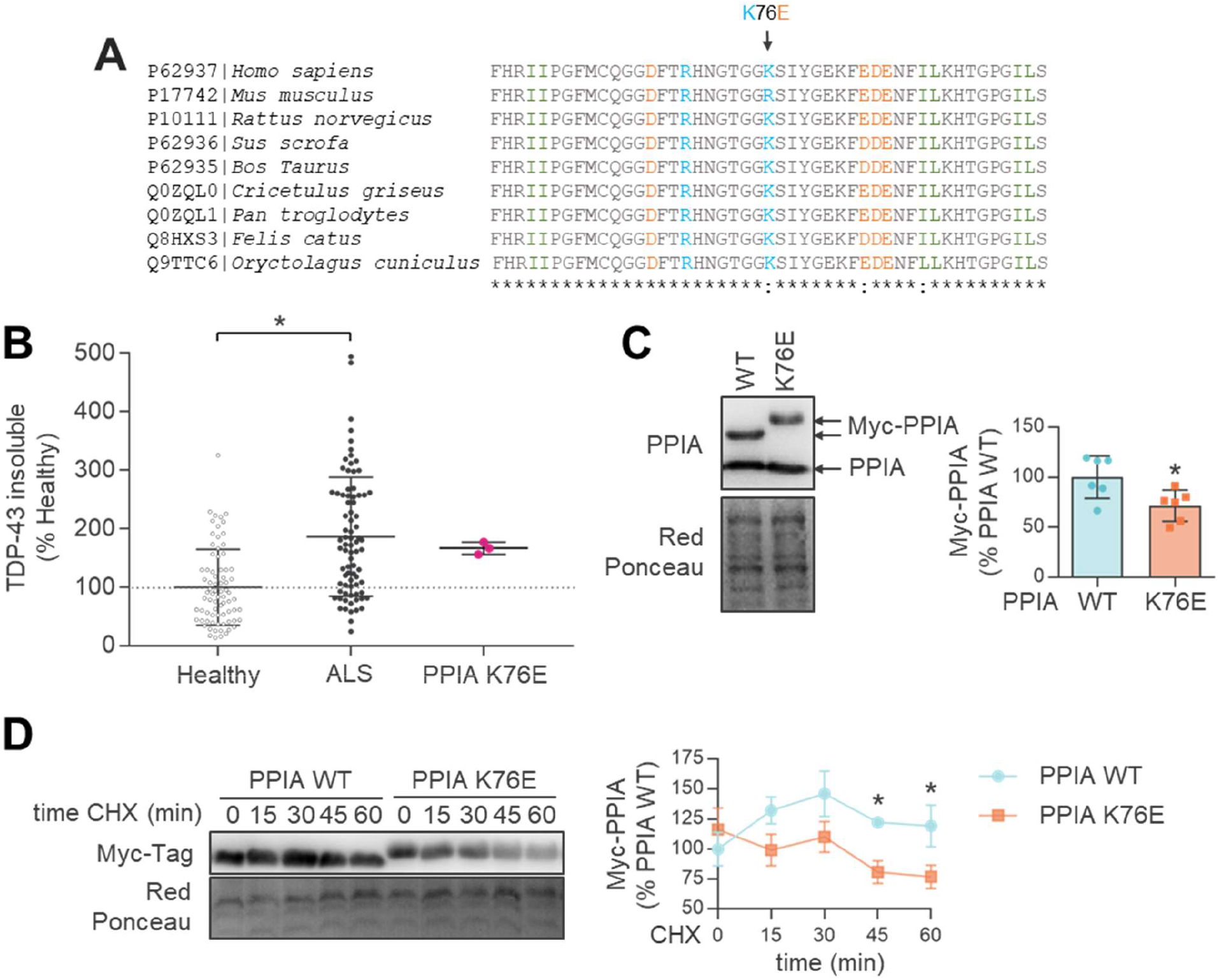
PPIA K76E mutant identified in a sporadic ALS patient. **(A)** PIA is evolutionarily highly conserved, 89% of the amino acid residues are identical in all the species listed. Multiple sequence alignment of PPIA, focused on the mutated lysine in position 76 and surrounding residues, is shown. (*), indicates conserved sites; (:) indicates conservative replacements (amino acids with similar biochemical properties); acidic residues (D, E) are in orange, basic residues (K, R) in blue, small aliphatic residues (I, L) in green, all other residues in grey. **(B)** Slot blot analysis of TDP-43 insoluble protein levels was done in PBMCs from the PPIA K76E ALS patient and compared to TDP-43 insoluble levels from ALS patients and healthy subjects of a retrospective cohort (Cohort #1, Supplementary Table 1). Immunoreactivity of the PPIA K76E patient was normalized to total protein loading and to the internal standard of the retrospective cohort ^7^, in order to compare the two analyses. Scatter dot plot (mean ± SD; n=76 healthy controls, n=79 ALS patients and n=3 technical replicates of the PPIA K76E ALS patient) are percentages of healthy controls and dotted line indicates the median value of healthy controls. *p < 0.05 versus healthy controls, Student’s t-test. **(C)** PPIA protein level was lower in HEK293 cells transfected with PPIA K76E mutant than PPIA WT plasmid, after 48 hours in culture. Immunoreactivity was normalized to protein loading. Representative WB are shown. Data (mean ± SEM; n=6) are percentages of PPIA WT. *p <0.05 versus PPIA WT Student’s t-test. **(D)** PPIA protein level was lower in HEK293 cells transfected with PPIA K76E mutant than PPIA WT plasmid after treatment with 100µg/mL cycloheximide (CHX) for 0, 15, 30, 45 and 60 minutes. Immunoreactivity was normalized to protein loading. Representative WB are shown. Data (mean ± SD; n=4) are percentages of PPIA WT. *p < 0.05 versus PPIA WT Student’s t-test.

We analysed TDP-43 partitioning in PBMCs as a surrogate marker of TDP-43 pathology and found that in the mutant PPIA patient TDP-43 accumulated in the insoluble fraction like the sporadic ALS and ALS-FTD patients of the retrospective cohort (Fig. 6B), confirming the diagnostic value of this parameter in PBMCs^7^. TDP-43 mRNA levels in the PPIA mutant patient were higher than in healthy controls and other ALS patients, while the protein level tended to be lower than the retrospective cohort (Supplementary Fig. 5C-D).

Known PPIA variants have low protein stability and are rapidly degraded^43^. We hypothesized that PPIA K76E could be a loss-of-function variant. We transfected HEK293 cells with myc-tagged PPIA wild-type (WT) and K76E and detected a lower level of the exogenous mutant protein than in WT after 48 hours in culture (Fig. 6C), despite equivalent transfection as detected in parallel wells by WB with anti-myc antibody after 24 hours in culture (Fig. 6D, timepoint 0). The lower stability of the mutant protein was confirmed by the accelerated degradation over time in cells treated with an inhibitor of new protein synthesis (Fig. 6D). Moreover, PPIA total protein levels in PBMCs were lower in the PPIA mutant patient than in the retrospective cohort^7^ of healthy subjects (−34%) and ALS patients (−21%) (Supplementary Fig. 5A). PPIA mRNA levels, however, were higher in the mutant PPIA patient than in other ALS patients, although lower than in healthy controls (Supplementary Fig. 5B).

We also examined possible structural differences between PPIA WT and K76E by molecular dynamics (MD) simulations (Fig. 7 and Supplementary Fig. 6). The analysis identified two regions (residues 26-30 and 43-45) outlining α-helix 1 in which the conformation of PPIA K76E significantly differed from the WT form (Fig. 7A-B). The mutant protein the 43-45 loop is significantly closer to the nearby β-strand (residues 156-165) (Fig. 7C). Since the K76E mutation lies within a coil in direct contact with helix-1, this suggested that the effect of the mutation is not only limited to a local change of electrostatic properties, but could instead result in a structural alteration of nearby regions essential for catalytic activity and protein interaction^44^. PPIA has a wide interaction network^13^. We tested whether the mutation impaired PPIA interaction with TDP-43 and Ran by co-immunoprecipitation (Fig. 7D-E). We found that both TDP-43 and Ran have an aberrant interaction with PPIA K76E, with a substantial increase in the affinity of both proteins for the mutant PPIA compared to PPIA WT.

**Figure 7.**
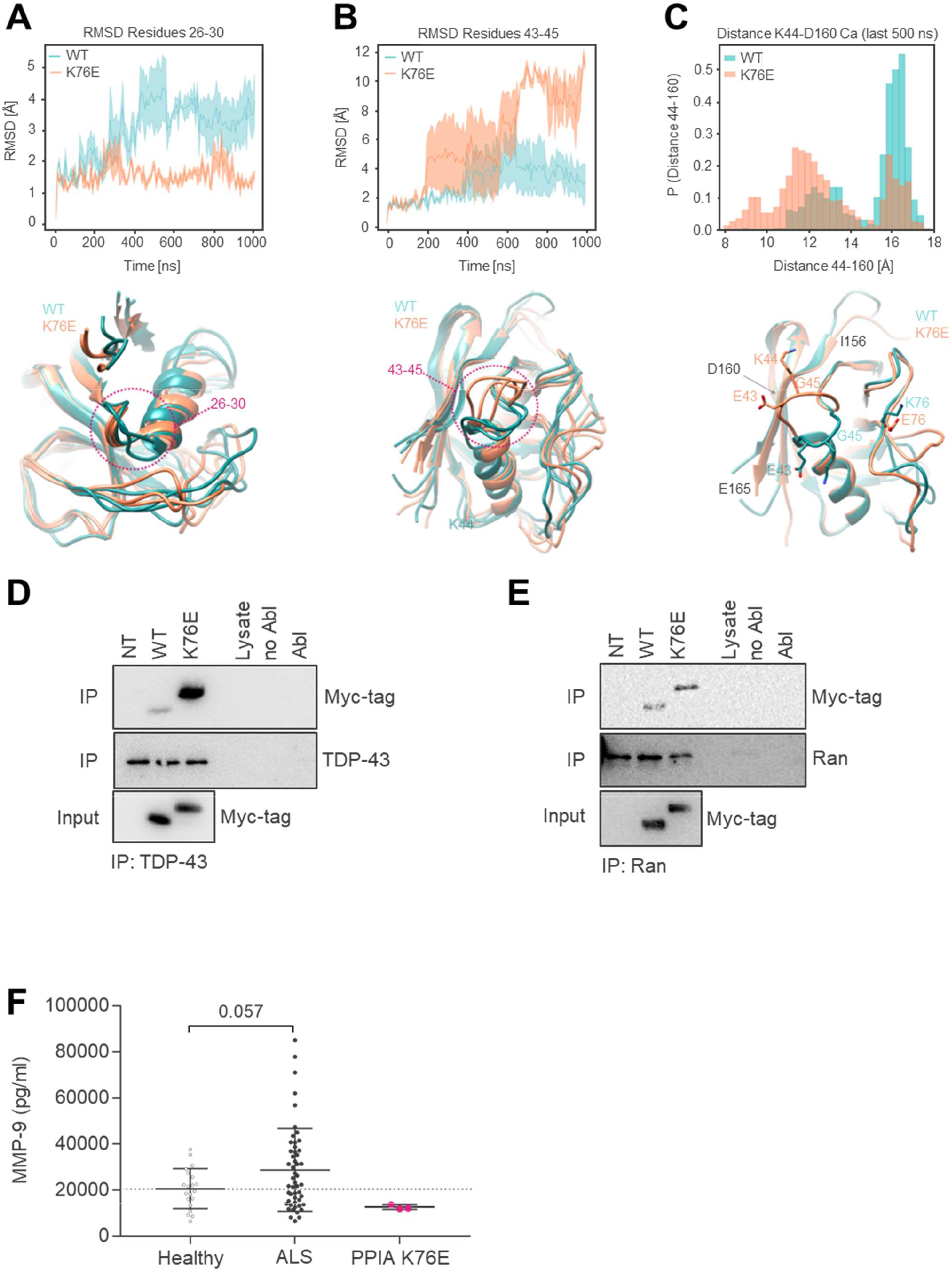
PPIA K76E shows structural differences from WT and is a loss-of-function mutant. **(A,B)** Structural analysis of protein loops composed residues 26-30 **(A)** and residues 43-45 **(B)** by molecular dynamics simulation. The protein RMSD of each frame, computed in comparison to the initial conformation, is plotted as a function of the simulation time (WT blue, K76E orange). Lines and filled curves represent the mean and standard error of the RMSD, respectively, computed for the three replicates of the WT and mutant protein. A significant difference between the two variants in loop 26-30 is evident after 400 ns of simulations **(A)**; in the loop 43-45 the difference is clear after 700 ns **(B)**. Three MD snapshots sampled at the end of the simulations for the WT protein (blue) superimposed on three MD snapshots sampled at the end of the simulations for the K76E variant (orange) are presented below the relative graph. **(C)** A detail of the structural variation of loop 43-45, with the probability distribution, computed considering the last 500 ns of dynamics, of the Cα distance between the residue at the center of the 43-45 loop (lysine 44) and the residue in the center of the β-strand 156-165 (aspartate 160). In the simulations of the K76E variant, the loop spent significantly more time closer to the β-strand 156-165 than the WT. The side-chains of the loop-residues face opposite directions in the two protein variants, as shown with representative conformations sampled after 1 μs of MD and superimposed below the graph. Here side-chains of residues E43, K44, G45 and K/E76 are explicitly represented (WT blue, K76 orange), while the positions of residues I156, D160 and E165 are indicated in grey. **(D-E)** Cells were transfected with PPIA WT or PPIA K76E construct, or not transfected (NT). TDP-43 **(D)** or Ran **(E)** was co-immunoprecipitated from cells using respectively an anti-TDP-43 polyclonal antibody or anti-Ran polyclonal antibody followed by anti-Myc WB. Controls for IP experiments: Lysate, lysate with magnetic beads linked to the secondary antibody (beads-IgGII); no AbI, lysis buffer with beads-IgGII; AbI, lysis buffer with beads-IgGII and primary antibody. WB anti-TDP-43 and anti-Ran show an equal amount of immunoprecipitated TDP-43 or Ran in PPIA WT, PPIA K76, NT cells. Representative WB are shown. **(F)** AlphaLISA analysis of MMP-9 was done in plasma samples from the PPIA K76E ALS patient, ALS patients and healthy subjects (Cohort #3, Supplementary Table 1). Scatter dot plots are mean ± SD (n=20 healthy controls, n=51 ALS patients and n=3 technical replicates of the PPIA K76E ALS patient). The dotted line indicates the median of healthy controls; p=0.057 versus healthy controls, Student’s t-test.

The extracellular form of PPIA is toxic for motor neurons by activating the EMMPRIN/CD147 receptor and inducing MMP-9 expression^18^. We measured MMP-9 plasma levels in the mutant PPIA patient and in a cohort of ALS patients and healthy subjects (Fig. 7F). While ALS patients had a higher level of MMP-9 than healthy controls, the PPIA mutant patient had one of the lowest levels of MMP-9, suggesting also a reduced toxic extracellular function of PPIA.

We conclude that K76E is a loss-of-function mutation that may affect both intracellular protective and extracellular toxic PPIA functions, leading to ALS with a slowly progressive phenotype in the patient. Further studies directly testing the function of mutant PPIA are necessary to dissect all possible implications.

## Discussion

TDP-43 proteinopathy is a prominent neuropathological feature in the ALS/FTD disease spectrum that is also seen in a wide range of neurodegenerative diseases, inherited and sporadic^4^. Neither the factors driving TDP-43 pathology nor its involvement in neurodegeneration have been defined yet. We propose that defective PPIA is a key factor in the cytoplasmic mislocalization and aggregation of TDP-43, induces neurodegeneration, and is a common feature in ALS/FTD patients.

PPIA is an evolutionarily conserved foldase and molecular chaperone, abundant in the cytoplasm but also present in the nucleus^7, 12^. Its PPIase activity has been regarded as fundamental in protein folding and refolding especially under stress conditions^13–15^. PPIA was found entrapped in aggregates isolated from spinal cord and brain of ALS/FTD patients and mouse models while exerting its protective function against misfolding^20, 45^. However, the absence of an overt phenotype in the *PPIA-/-* mouse in the work of Colgan et al. and its dispensable function in mammalian cells have kept its physiological role enigmatic^24, 25^, especially in the CNS where it is highly expressed.

In a previous work characterizing the protein interacting network of PPIA under basal conditions we found that it includes several proteins involved in RNA homeostasis, hnRNPs and TDP-43^13^. We also reported that PPIA regulates key TDP-43 functions, such as gene expression regulation^13^. In fact, knocking down PPIA affected the expression of TDP-43 RNA targets implicated in FTD/ALS, such as GRN, VCP and FUS, or in protein clearance, such as HDAC6 and ATG7. As a consequence, the mutant SOD1 mouse cross-bred with the *PPIA-/-* mouse had a more severe disease phenotype, with more protein aggregation and less survival. Besides SOD1, TDP-43, and hnRNPs^13^, PPIA interacts with other proteins linked to genetic forms of ALS/FTD, such as ubiquilin 2^46^ and Arg-containing dipeptide repeat proteins^47^. We therefore concluded that PPIA could be a modifier of ALS/FTD pathology.

In this work we demonstrate that the loss of PPIA in mice is sufficient to induce neurodegeneration leading to a clinical phenotype of the ALS/FTD spectrum. We also show that PPIA is defective in several ALS and ALS-FTD patients and that a subject carrying a loss-of-function PPIA mutation develops ALS. Therefore, besides the general intracellular protective effect under stress, we provide evidence that PPIA has specific, non-redundant functions that are particularly relevant for TDP-43 biology.

Deficient PPIA alters the biochemical properties of Ran and this may underlie the impaired TDP-43 nuclear import. Ran is a master regulator of the nuclear protein import and is required for the majority of the proteins that shuttle between the nucleus and cytoplasm. Non-functional Ran reduced TDP-43 nuclear localization in cortical neurons^38^. Ran is also a PPIA interactor^13^ and has a GP-motif (residues 57-58), the recognition site for the foldase activity of PPIA, present also in TDP-43 and other PPIA substrates ^13, 44, 48, 49^. In the absence of PPIA, Ran is increasingly recovered in the detergent-insoluble aggregates of the mouse brain. Ran and other regulators of nucleocytoplasmic trafficking accumulate in the cytoplasm of ALS/FTD cellular models^50–52^, either as de-mixed droplets or in stress granules, suggesting that defective PPIA may have a role in the pathogenic mechanisms that couple cellular stress with impaired nucleocytoplasmic transport. Biochemical analysis of the detergent-insoluble fraction of the cortex of *PPIA-/-* mice did in fact indicate a general increase in insoluble proteins and TIA1 stress granule marker.

Defective PPIA alters TDP-43 mRNA expression. In *PPIA-/-* mice TDP-43 mRNA is downregulated, and in the mutant PPIA patient it is upregulated. TDP-43 physiological levels are tightly controlled at a transcriptional level by an autoregulatory loop, which keeps intracellular TDP-43 within a narrow range^2, 3^. This is necessary, for example, for TDP-43’s role in RNP complexes that may become dysfunctional if the stoichiometry between TDP-43 and the other protein/RNA components is disrupted^13^. In a previous work, we demonstrated that PPIA through its foldase activity influences the binding of TDP-43 to RNA^13^. Therefore, we can assume that in absence of *PPIA-/-* increased *TARDBP* intron 7 splicing and TDP-43 mRNA destabilization in mice may be caused by an impaired binding of TDP-43 to its 3′ UTR. In the mutant patient, we can hypothesize that PPIA K76E, that has an aberrant interaction with TDP-43, sequesters TDP-43 and affects autoregulation in the opposite direction, promoting the production of stable mRNA species^53^. Further studies are needed to decipher the exact mechanism at the basis of these divergent effects at a mRNA level, while at a protein level in both cases TDP-43 decreases. However, TDP-43 homeostasis and function are controlled not only by autoregulation, but also by nucleocytoplasmic transport and aggregation, in a coordinated fashion^10^. Interestingly, PPIA seems to be involved in all of them.

In brain of *PPIA-/-* mice TDP-43 is downregulated also at the protein level. If complete ablation of TDP-43 is embryonically lethal, conditional knock-out and limited knock-down of TDP-43 results in neurodegeneration^54, 55^. Moreover, TDP-43 depletion in brain downregulates genes with very long introns that encode proteins involved in synaptic function^3, 56^. In agreement with this, we detected downregulation of synaptic proteins and impairment of synaptic plasticity in the hippocampus of *PPIA-/-* mice, indicating that PPIA deficiency also has an effect on synaptic structure and function.

In several pathological conditions oxidative stress and inflammation increase PPIA secretion into biofluids^57^. The levels of PPIA were high in the CSF of ALS patients and mutant SOD1 rodent models^18^. Once secreted, PPIA activates a proinflammatory pathway by the EMMPRIN/CD147 receptor, leading to NF-κB activation and induction of MMP-9 expression in neurons and glia^18^. Large, fast-twitch, fatigable α motor neurons, which are the most susceptible to degeneration in ALS, express the highest level of MMP-9, which enhances endoplasmic reticulum stress and promotes neurodegeneration^58^. High levels of MMP-9 in tissues and biofluids of ALS patients have been detected in this work and by other groups^59, 60^ and has been associated with blood-brain barrier breakdown^61^. We previously reported that selective pharmacological inhibition of extracellular PPIA (ePPIA) reduced MMP-9, promoted a pro-healing phenotype in glia, rescued motor neurons and increased survival in the SOD1 G93A mouse model of ALS^18^. We also showed that ePPIA was toxic toward motor neurons, which express high levels of EMMPRIN/CD147, but had no effect on cortical and cerebellar granule neurons^18^.

The late-onset and slowly progressing motor dysfunction in *PPIA-/-* mice can be explained with the substantially reduced activation of the EMMPRIN/CD147-dependent proinflammatory pathway to which motor neurons are particularly susceptible (Fig. 8). In the patient heterozygous for mutant PPIA, half the ePPIA may be unable to activate EMMPRIN/CD147, reducing neuroinflammation and slowing progression of the disease. Defective intracellular PPIA (iPPIA) increases cellular stress, ultimately leading to PPIA secretion. We hypothesize a tug-of-war between iPPIA and ePPIA, in which the prevalence of ePPIA over iPPIA accelerates disease progression. Increasing iPPIA function may reduce ePPIA and be an effective therapeutic approach.

**Figure 8.**
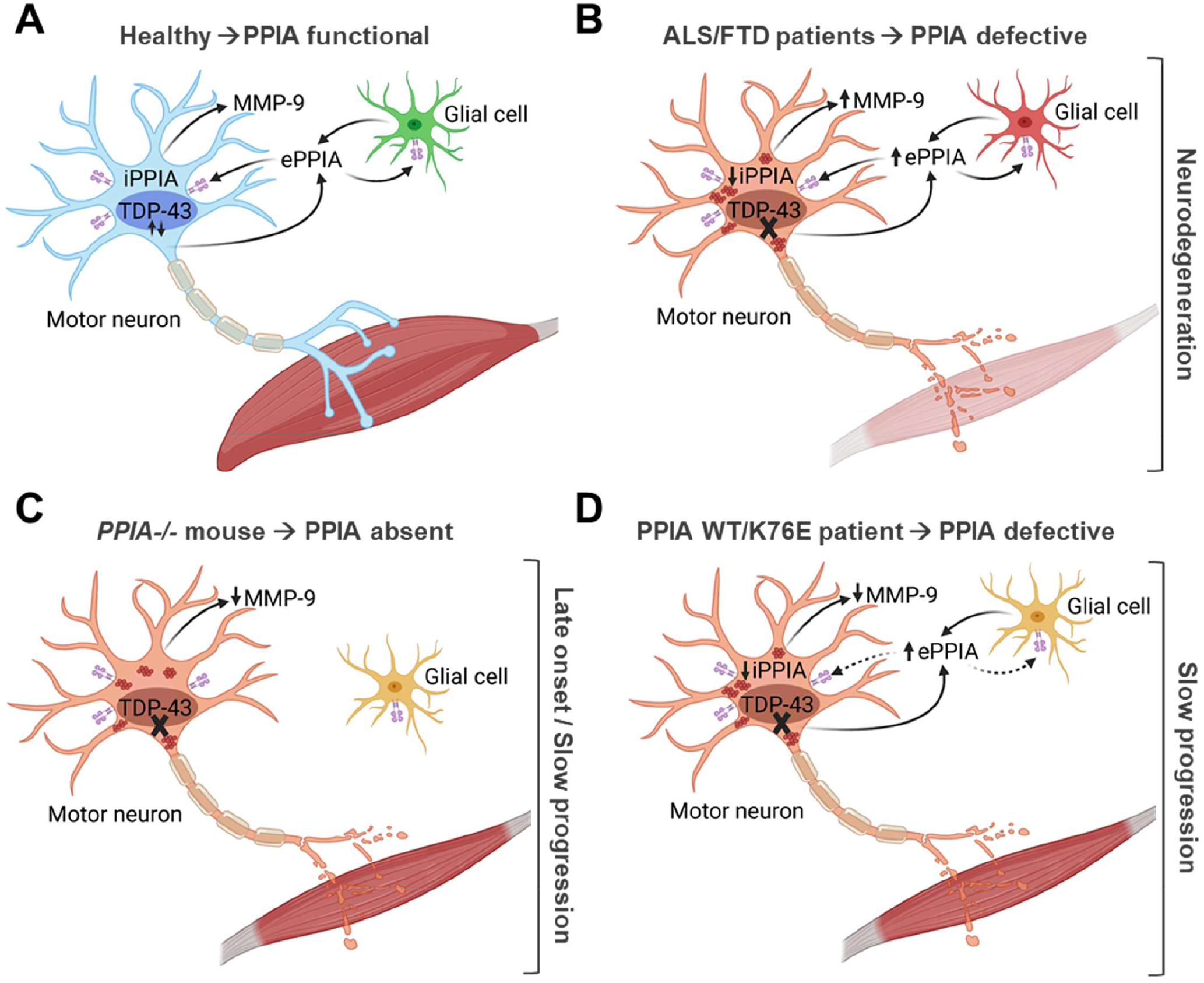
Schematic representation of PPIA function in physiological and pathological conditions. **(A)** PPIA (iPPIA) is highly expressed in neurons and motor neurons where it is protective thanks to its activity as a foldase and molecular chaperone that affects key proteins involved in RNA metabolism, nucleocytoplasmic transport and synaptic function. PPIA is secreted (ePPIA) in response to oxidative stress and inflammatory stimuli^57^. **(B)** In ALS/FTD, defective iPPIA induces TDP-43 pathology, nucleocytoplasmic transport defects, synaptic dysfunction and neurodegeneration. Persistent cellular stress in neurons and glia leads to aberrant secretion of PPIA that, through the interaction with its EMMPRIN/CD147 receptor, induces MMP-9 expression and an inflammatory response which is particularly toxic for motor neurons^18, 58^. **(C,D)** In the *PPIA-/-* mouse **(C)** and PPIA WT/K76E patient **(D)**, the detrimental effect of deficient iPPIA function is partially compensated by the lack (mouse) or lower (patient) ePPIA-induced inflammatory response, leading to marked TDP-43 pathology associated with late onset/slowly progressive (mouse) or slowly progressive (patient) motor neuron disease.

Several mouse models of ALS/FTD have been generated targeting genes with known pathogenic roles but each one recapitulates only certain aspects of the human disease. For instance, TDP-43 pathology in mice is often absent or mild and is not always linked to neurodegeneration or behavioural phenotypes^62^. In TDP-43 mouse models overexpressing wild-type TDP-43 or disease-associated mutations, C-terminal TDP-43 fragments are detected at low levels or are absent^63^. *PPIA-/-* mice are characterized by diffuse, marked TDP-43 pathology in cortex, hippocampus and spinal cord^13^, with all the key neuropathological features of ALS/FTD including cytoplasmic mislocalization and accumulation of C-terminal fragments of TDP-43 associated with neurodegeneration and cognitive and behavioural impairments. In *PPIA-/-* mice the behavioural symptoms follow a peculiar course. Initially, they present disinhibition with no social impairment, and later they show loss of disinhibition and develop apathy and social disinterest. Interestingly, while most patients with the behavioural variant of FTD (bvFTD) display both disinhibition and apathy during the course of the disease, some may initially present as primarily disinhibited or primarily apathetic and later develop either signs of inertia or disinhibition^64^. Moreover, despite clear evidence that cognitive/behavioural impairment may appear early in the course of ALS not always deterioration seems to occur with disease progression^65^.

There is a significant genetic and neuropathological overlap between ALS and FTD with up to 50% of ALS patients having cognitive and behavioural manifestations and up to 40% of FTD patients developing motor dysfunction^66^. However, it is not clear which pathway is at the basis of either phenotype. Interestingly, *PPIA-/-* mice display early key symptoms of bvFTD and only later develop motor dysfunction. The resulting picture resembles a behaviourally predominant ALS-FTD in which behavioural symptoms typically evolve before motor symptoms^67^. GRN deficiency, the cause of a tau-negative familial form of FTD^68^, probably contributes to the development of a predominant FTD phenotype in *PPIA-/-* mice.

PPIA genetic variants are very rare, so individuals homozygous for any of them even more unlikely^43^. The identification of a PPIA variant in a single patient cannot definitely demonstrate its causality. However, the structural and functional properties of the mutant protein are suggestive of its possible involvement in disease pathogenesis. More relevant for the disease is the regulation of PPIA at a post-translational level. PPIA is commonly and variably post-translationally modified and these modifications regulate its functions^13, 21, 69, 70^. The interaction with TDP-43 is favoured by PPIA Lys-acetylation, which is low in ALS patients^13^. Here we also report downregulated PPIA in two independent cohorts of patients; therefore, we predict that an overall defective PPIA is likely in several patients and needs to be further investigated in large clinical studies.

In conclusion, our findings indicate that PPIA is involved in multiple pathways that protect CNS from TDP-43-mediated toxicity. In fact, if deficient and/or dysfunctional PPIA can trigger TDP-43 proteinopathy leading to neurodegeneration. *PPIA-/-* mice recapitulate key features of ALS-FTD and are useful experimental model to investigate the mechanisms of TDP-43 proteinopathy, with the aim of developing novel therapeutic approaches.

## Acknowledgments

We thank Bradford C. Berk and Patrizia Nigro for providing the *PPIA-/-* mice on C57BL/6J genetic background and the Laboratory of Neurogenetics (NIH) staff for their collegial support and technical assistance. We thank Judith Baggott for editorial assistance and Fabrizio Aspesi for graphics assistance by BioRender.com.

## Supplementary Material

-Materials and methods

-Supplementary Table 1

-Supplementary figure legends

-Supplementary Fig. 1

-Supplementary Fig. 2

-Supplementary Fig. 3

-Supplementary Fig. 4

-Supplementary Fig. 5

-Supplementary Fig. 6

## Materials and methods

### Animal model

Animals were bred and maintained at the Istituto di Ricerche Farmacologiche Mario Negri IRCCS, Milan, Italy, under standard conditions: temperature 21 ± 1°C, relative humidity 55 ± 10%, 12h light schedule, and food and water ad libitum. Before every analysis, animals were deeply anesthetized with ketamine hydrochloride (IMALGENE, 100 mg/kg; Alcyon Italia) and medetomidine hydrochloride (DOMITOR, 1 mg/kg; Alcyon Italia) by intraperitoneal injection and killed by decapitation. *PPIA-/-* mice were originally generated and characterized as described^24, 25^. We obtained *PPIA-/-* mice (strain 129S6/SvEvTac Ppiatm1Lubn/Ppiatm1Lbn; stock no. 005320) from The Jackson Laboratory; they were maintained on a 129S6/SvEvTac background. The *PPIA-/-* mice on 129S6/Sv genetic background and corresponding *PPIA+/+* littermates were used for micro-CT, MRI, immunohistochemistry, biochemistry, long-term potentiation, Rotarod, grid test, extension reflex and are indicated as *PPIA-/-* mice. *PPIA-/-* mice on C57BL/6J genetic background (C57_*PPIA-/-* mice), kindly provided by Dr. Bradford C. Berk (University of Rochester Medical Center, Rochester, New York, USA), and corresponding *PPIA+/+* littermates, were used for open field, elevated-plus maze, three-chamber sociability, Morris water maze, novel object recognition. We verified that C57_*PPIA-/-* mice also display TDP-43 pathology (Supplementary Fig. 3). Genotyping for PPIA was done by standard PCR on DNA tail biopsies, using primer sets designed by The Jackson Laboratory. The number of animals was calculated on the basis of experiments designed to reach a power of 0.8, with a minimum difference of 20% (α = 0.05).

### Micro-CT

Micro-CT was done with an Explore Locus micro-CT scanner (GE Healthcare) without contrast agents. Before the analysis, mice were anesthetized with a continuous flow of 3% isofluorane/oxygen mixture and placed prone on the micro-CT bed. A micro-CT lower resolution (Bin-4) protocol was employed using 80 kV, 450 μA with 100 msec per projection and 400 projections over 360° for a total scan time of 10 minutes, as previously described^27^. The isotropic resolution of this protocol is 93 μm. The scanned images were reconstructed in 3D and analysed using Micro View analysis software (version 2.1.1; GE Healthcare). We measured the distance from the last cervical vertebra to the last lumbar vertebra (segment AB; Supplementary Fig. 1C) and the perpendicular distance to the dorsal edge of the vertebra at the greatest curvature (segment CD; Supplementary Fig. 1C). The index of kyphosis was defined as the ratio of AB to CD.

### MRI analysis

MRI analysis was done in six- and twelve-month-old mice, as previously described ^26^. Briefly animals were anesthetized with isoflurane in a mixture of O_2_ (30%) and N_2_O (70%). Body temperature was maintained at ∼37°C by a warm-water-circulated heating cradle. Imaging was done on a 7 T small bore animal Scanner (Bruker Biospec, Ettlingen, Germany). A 3D RARE T2 weighted sequence was used to assess anatomical changes. The volume measurements of structural MRI images were obtained using Java-based custom-made software. ROIs were selected manually and drawn on the images for volumetric assessment. Whole brain, hippocampus, cortex and cerebellum were measured. Cortex was quantified from the rhinal fissure (RF) up. Data from each animal were obtained by integrating the averaged ROI area for slice thickness, and normalized to whole brain volume.

### Immunohistochemistry

Mice were anesthetized and perfused transcardially with 50 mL of phosphate-buffered saline (PBS) followed by 100 mL of 4% paraformaldehyde (Sigma-Aldrich) in PBS. Brain and spinal cord were rapidly removed, postfixed for 3h, transferred to 20% sucrose in PBS overnight and then to 30% sucrose solution until they sank, frozen in N-pentane at 45°C and stored at ± 80°C. Before freezing, spinal cord was divided into cervical, thoracic, and lumbar segments and included in Tissue-tec OCT compound (Sakura). Coronal sections (30 μm; four slices per mouse) of brain were then sliced and immunohistochemistry was done for TDP-43, phosphorylated TDP-43 (pTDP-43), glial fibrillary acidic protein (GFAP) and Iba-1. Briefly, slices were incubated for 1h at room temperature with blocking solutions (TDP-43 and pTDP-43: 0.2% Triton X100 plus 2% normal goat serum (NGS); GFAP: 0.4% Triton X100 plus 3% NGS; Iba-1: 0.3% Triton X100 plus 10% NGS), then overnight at 4°C with the primary antibodies (TDP-43, 1:200; pTDP-43, 1:1000; GFAP, 1:2500; Iba-1, 1:1000). Brand and Research Resource Identifiers (RRID) of the antibodies are reported in the Immunoblotting section. After incubation with biotinylated secondary antibody (1:200; 1h at room temperature; Vector Laboratories) immunostaining was developed using the avidin–biotin kit (Vector Laboratories) and diaminobenzidine (Sigma). Coronal brain sections (30 μm; four slices per mouse) and lumbar spinal cord (30 μm; twelve slices per mouse) were stained with 0.5% cresyl violet to detect the Nissl substance of neuronal cells. Stained sections were collected at 20 X with an Olympus BX-61 Virtual Stage microscope so as to have complete stitching of the whole section, with a pixel size of 0.346 μm. Acquisition was done over 6-μm-thick stacks with a step size of 2 μm. The different focal planes were merged into a single stack by mean intensity projection to ensure consistent focus throughout the sample. Finally, signals were analysed for each slice with ImageJ software.

### Subcellular fractionation

Nuclear and cytoplasmic fractions were separated from mouse cortex and cerebellum as described ^13, 18^. Briefly, tissues were homogenized in six volumes (w/v) of buffer A (10 mM Tris-HCl pH 7.4, 5 mM MgCl2, 25 mM KCl, 0.25 M sucrose, 0.5 mM DTT) containing a protease inhibitors cocktail (Roche), and centrifuged at 800 xg for 10 min at 4°C. The supernatant was centrifuged twice at 800 xg for 10 min at 4°C (cytoplasmic fraction). The pellet was resuspended in three volumes of buffer A and centrifuged three times at 800 xg for 10 min at 4°C. The pellet was resuspended in one volume of buffer A and one volume of buffer B (10 mM Tris-HCl pH 7.4, 5 mM MgCl2, 25 mM KCl, 2 M sucrose) containing a protease inhibitors cocktail, and loaded on a layer of one volume of buffer B. Samples were ultracentrifuged at 100,000 xg for 45 min at 4°C. The pellet (nuclear fraction) was resuspended in 100 μL of buffer A, centrifuged at 800 xg for 10 min at 4°C and resuspended in 40 μL buffer A.

### Extraction of detergent-insoluble

Mouse tissues were homogenized in 10 volumes (w/v) of buffer, 15 mM Tris-HCl pH 7.6, 1 mM DTT, 0.25 M sucrose, 1 mM MgCl_2_, 2.5 mM EDTA, 1 mM EGTA, 0.25 M sodium orthovanadate, 2 mM sodium pyrophosphate, 25 mM NaF, 5 µM MG132, and a protease inhibitors cocktail (Roche), essentially as described ^13^. Samples were centrifuged at 10,000 xg at 4°C for 15 min and supernatant 1 was collected in a new tube. The pellet was suspended in ice-cold homogenization buffer with 2% Triton X100 and 150 mM KCl, sonicated and shaken for 1h at 4°C. The samples were then centrifuged twice at 10,000 xg at 4°C for 10 min to obtain the Triton-insoluble fraction (TIF) pellet and supernatant 2. Supernatants 1 and 2 were pooled, as the Triton-soluble fraction. Immunoreactivity was normalized to protein loading (Ponceau red staining). The amount of Triton-resistant proteins isolated from the tissue was normalized to the soluble protein extracted. Proteins were quantified by the BCA protein assay (Pierce).

### Immunoblotting

Protein levels were determined using the BCA protein assay (Pierce). For western blot (WB), samples (15 µg) were separated in 12% SDS-polyacrylamide gels and transferred to polyvinylidene difluoride membranes (Millipore), as described previously ^20^. For dot blot, proteins (3 µg) were loaded directly onto nitrocellulose Trans-blot transfer membranes (0.45 µm; Bio-Rad), depositing each sample on the membrane by vacuum filtration, as described previously ^7^. WB and dot blot membranes were blocked with 3% (w/v) BSA (Sigma-Aldrich) and 0.1% (v/v) Tween 20 in Tris-buffered saline, pH 7.5, and incubated with primary antibodies and then with peroxidase-conjugated secondary antibodies (GE Healthcare). Antibodies used for immunoblot are the following: rabbit polyclonal anti-PPIA antibody (1:2500, Proteintech; RRID: AB_2237516); rabbit polyclonal anti-human C-terminal TDP-43 antibody (1:2500, Proteintech; RRID: AB_2200505); mouse monoclonal anti-human pTDP-43 antibody (1:2000, Cosmo Bio Co., LTD; RRID: AB_1961900) ^71^; rabbit polyclonal anti-Ran antibody (1:1000, Cell Signaling; RRID: AB_2284873); mouse monoclonal anti-PSD95 antibody (1:10000, Neuromab; RRID: AB_2292909); rabbit polyclonal anti-synaptophysin antibody (1:5000, Synaptic System; RRID: AB_887905); mouse monoclonal anti-Myc Tag antibody (1:1000, Millipore; RRID: AB_11211891); mouse monoclonal anti-GFAP antibody (1:1000; Millipore; RRID:AB_94844); rabbit polyclonal anti-Iba-1 antibody (1:500; Wako; RRID:AB_839504); mouse monoclonal anti-hnRNPA2/B1 antibody (1:2000, Abnova; RRID: AB_425488); rabbit polyclonal anti-TIA1 antibody (1:1000, Proteintech; RRID: AB_2201427); goat anti-mouse or anti-rabbit peroxidase-conjugated secondary antibodies (respectively 1:20000 and 1:10000, GE Healthcare). Blots were developed with the Luminata Forte Western Chemiluminescent HRP Substrate (Millipore) on the ChemiDoc™ Imaging System (Bio-Rad). Densitometry was done with Progenesis PG240 version 2006 software (Nonlinear Dynamics) and Image Lab 6.0 software (Bio-Rad). The immune reactivity of the different proteins was normalized to Ponceau Red staining (Fluka).

### Real-time PCR

The total RNA from mouse cortex and human PBMC was extracted using the RNeasy® Mini Kit (Qiagen). RNA samples were treated with DNase I and reverse transcription was done with a High-Capacity cDNA Reverse Transcription Kit (Life Technologies). For human and mouse PPIA, *TARDBP* and mouse GRN real-time PCR we used the Taq Man Gene expression assay (Applied Biosystems), on cDNA specimens in triplicate, using 1X Universal PCR master mix (Life Technologies) and 1X mix containing specific receptor probes for mouse GRN (Mm00433848_m1; Life Technologies), mouse *TARDBP* (Mm01257504_g1; Life Technologies), human PPIA (Hs03045993_gH; Life Technologies) and human *TARDBP* (Hs00606522_m1; Life Technologies). Relative quantification was calculated from the ratio of the cycle number (Ct) at which the signal crossed a threshold set within the logarithmic phase of the given gene to that of the reference mouse β-actin gene (Mm02619580_g1; Life Technologies) or human β-actin gene (Hs01060665_g1; Life Technologies). For *TARDBP* intron 7 exclusion real time PCR we used SYBR Green assay (Applied Biosystem), on cDNA specimens in triplicate, using 1X SYBR Green (Applied Biosystem) and specific primers for mouse *TARDBP* Intron 7 (F: TTCATCTCATTTCAAATGTTTATGGAAG; R: ATTAACTGCTATGAATTCTTTGCATTCAG)^53^. Relative quantification was calculated from the ratio of the cycle number (Ct) at which the signal crossed a threshold set within the logarithmic phase of the given gene to that of the reference mouse β-actin gene (F: GCCCTGAGGCTCTTTTCCAG R: TGCCACAGGATTCCATACCC). For both experiments means of the triplicate results for each sample were used as individual data for 2^-ΔΔCt^ statistical analysis.

### Long-term potentiation analysis

For extracellular recordings coronal brain slices (350 µm) were cut in ice-cold modified artificial cerebrospinal fluid (aCSF) containing the following: 87 mM NaCl, 2.5 mM KCl, 1 mM NaH_2_PO_4_, 75 mM sucrose, 7 mM MgCl_2_, 24 mM NaHCO_3_, 11 mM D-glucose, and 0.5 mM CaCl_2_. Slices where then transferred to an incubating chamber, submerged in aCSF containing 130 mM NaCl, 3.5 mM KCl, 1.2 mM NaH_2_PO_4_, 1.3 mM MgCl_2_, 25 mM NaHCO_3_, 11 mM D-glucose, 2 mM CaCl_2_, constantly bubbled with 95% O_2_ and 5% CO_2_ at room temperature. Slices were incubated in this condition for at least 1h before recording, then transferred into a submerged recording chamber and perfused with oxygenated aCSF at 2 mL/min at a constant temperature of 28-30°C. Field EPSPs (fEPSPs) were recorded with glass micropipettes filled with 3 M NaCl electrode in CA1 stratum radiatum. The Schaffer collaterals were stimulated with a bipolar twisted Ni/Cr stimulating electrode. Stimuli were delivered by a Constant Voltage Isolated Stimulator (Digitimer Ltd., Welwyn Garden City, UK). Data were amplified and filtered (10Hz to 3kHz) with a DAM 80 AC Differential Amplifier (World Precision Instruments, Sarasota, FL), and digitized at 10 kHz by a Digidata 1322 (Molecular Devices, Foster City, CA). Long-term potentiation (LTP) was induced by a 4-theta burst tetanus stimulation protocol (each burst consists of four pulses at 100 Hz with 200 ms inter-burst intervals). LTP recordings in which the amplitude of the presynaptic fiber volley changed by more than 20% were discarded.

### Behavioural analysis

In this study both male and female mice were tested, at 6 and 12 months of age. All behavioural tests were done at the same time of day, in the afternoon. Mice were allowed to habituate to the test room for at least 1h. Test environments were thoroughly cleaned between test sessions and males were tested before females. Mice were weighed once every 15 days, at the same time of day, in the afternoon, on a balance with 0.1 g readability and ± 0.3 g linearity (EU-C 7500PT, Gibertini). The open field, elevated-plus maze, three-chamber sociability, Morris water maze and novel object recognition test used Ethovision XT, 5.0 software (Noldus Information Technology, Wageningen, The Netherlands) to record the parameters. Mice were randomized and the operators were blinded for the behavioural analysis.

### Open field

The open field consists of a square Perspex box arena with walls (40 x 40 x 40 cm). The “center zone” was defined as a square covering 16% of the total arena area (16 x 16 cm central square) and the “periphery zone” as the surrounding border. Each mouse was placed in the center of the arena and velocity, total distance moved and the time spent in the central and periphery zones of the open field were recorded for 5 min. Velocity was analysed as a measure of the locomotor activity.

### Elevated-plus maze

The device consisted of a central part (5 x 5 cm), two opposing open arms (30 x 5 cm) and two opposing closed arms (same size) with 14 cm high, non-transparent walls. The maze was elevated 73 cm off the floor. At the beginning of each trial, mice were placed on the central platform, facing an open arm. Their behaviour was recorded for 5 min then analysed by an operator blind to the genotype. Entry into an arm was recorded when the mouse placed its four paws in that arm^72^.

### Three-chamber sociability

The apparatus for the social behaviour test consists of three chambers connected with retractable open doorways on two dividing walls. The two external chambers contained an inverted empty wire cup. The test comprised three main steps. After 5 min habituation, during which the test mouse was allowed to explore the arena freely, an unfamiliar mouse (stranger 1) was introduced into one of the empty wire cups to measure social preference. The time the test mouse spent sniffing each wire cup was recorded for 10 min, then a new mouse (stranger 2) was introduced into the other empty wire cup to measure social recognition. Time spent sniffing each wire cup was measured. Unfamiliar mice of the same sex and age were used as strangers 1 and 2.

### Morris water maze

The Morris water maze test (MWM) consists of a round pool (100 cm diameter) filled with water (22°C) made opaque by the addition of nontoxic, odorless white tempera. The escape platform, made of transparent plastic, was placed 1 cm below the water surface. The test involved five days of training and a one-day probe trial. During each day of training mice were placed successively in north, east, south, and west positions, with the escape platform hidden in the middle of the southwest quadrant. Whenever the mice failed to reach the escape platform, they were placed on it for 10 s. Latencies before reaching the platform were recorded in four-trial sessions. After the training, on the sixth day a probe trial was conducted, removing the platform from the pool. Time spent in the previously correct target quadrant (Q1) was measured in a single 1-min trial.

### Novel object recognition

The novel object recognition test was done in a square gray arena with walls (40 x 40 x 40 cm). The following objects were used: a black plastic cylinder (4 x 5 cm) and a metal cube (3 x 5 cm). After 5 min habituation during which the animal explored the empty arena freely, on the second day mouse was placed in the same arena containing two identical objects. On the last day mouse was again placed in the arena containing the object presented the second day (familiar), together with a new, different one, and the time spent exploring the two objects was recorded for 5 min. The parameter analysed was the discrimination index (DI), defined as: DI = (T_New_ – T_Familiar_) / (T_New_ + T_Familiar_) where T_New_ is the time spent with the new object and T_Familiar_ the time spent with the familiar object.

### Rotarod

Motor performance of the mice was determined using a Rotarod (Ugo Basile) in acceleration mode (7-44 rpm) over 5 min. The mice were allowed up to three attempts and the longest latency to fall was considered in statistical analysis. Mice were tested at the same time of day, in the afternoon, once every 15 days, from 13 to 22 months of age.

### Grid test

Grip strength was measured using the following score:

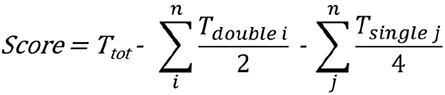

where T_tot_ is the time spent hanging before falling from the grid, n is the number of events in which both hind paws (i) or a fore or a hind paw (j) became detached from the grid, T_double i_ is the number of seconds the i-th event lasted, T_single j_ is the number of seconds the j-th event lasted, as previously described ^13^. Mice were tested at the same time of the day, in the afternoon, once every 15 days, from 13 to 22 months of age.

### Extension reflex

Extension reflex was quantified using the following 3-point score: 3, hindlimbs extended to an angle of 120 degrees; 2.5, hindlimbs extended to < 90 degrees with decreased reflex in one limb; 2.0, as 2.5 with decreased reflex in both hindlimbs; 1.5, loss of reflex with marked flexion in one hindlimb; 1, as 1.5 with marked flexion in both hindlimbs; 0.5, loss of reflex with hindlimbs and paws held close to the body. Mice were tested at the same time of day, in the afternoon, once every 15 days, from 13 to 22 months of age.

### Isolation of PBMCs and plasma

Blood was drawn by standard venipuncture into Vacutainer® Plus Plastic K2EDTA Tubes (Becton, Dickinson and Company) and kept at 4°C until shipment to the Istituto di Ricerche Farmacologiche Mario Negri IRCCS. PBMCs were isolated from ALS patients and healthy individuals and proteins were extracted as previously described ^7^. Briefly, PBMC were isolated from EDTA-blood by Ficoll-Hypaque (Ficoll-Plaque™ Plus, GE Healthcare) density gradient centrifugation. Mononuclear cells were harvested from the interface and washed with RPMI 1640 medium (EuroClone). To extract proteins, PBMC pellets were suspended with buffer 1 (20 mM Tris-HCl pH 7.5, 0.1% NP40, 0.1% SDS, cOmplete™ Protease Inhibitor Cocktail Tablets, Roche), kept at 95°C and shaken at 800 rpm for 5 min, then centrifuged at 16,000 xg for 10 min at 4°C. The supernatants (Soluble) were collected and the pellet (Insoluble) was suspended with buffer 2 (1% SDS at 95°C), kept at 95°C and shaken at 800 rpm for 5 min). To isolate plasma samples, blood was centrifuged at 3000 xg for 20 minutes and plasma was stored at -80°C until use.

### Mutation screening

We examined a cohort of 959 ALS patients from Northern Italy and 677 healthy controls matched by age, sex and geographical origin. Informed written consent was obtained for all subjects and the study was approved by the ethics committees involved. Standard protocols were followed for DNA extraction from peripheral blood. Whole-genome sequencing was done at The American Genome Center (Uniformed Services University, Walter Reed National Military Medical Center campus, Bethesda, MD, USA). Libraries were prepared using TruSeq DNA PCR-Free High Throughput Library Prep Kit (Illumina Inc.). Sequencing was done on an Illumina HiSeqX10 sequencer using paired-end 150 base pair reads. Sequence reads were processed as per Genome Analysis Toolkit’s (GATK) standard practices (https://software.broadinstitute.org/gatk/best-practices/). The GATK variant quality score method with default filters was used for variant quality control. Genome Reference Consortium Human Build 38 was used as the reference. Variant annotation was done using KGGseq version 1.0 (http://grass.cgs.hku.hk/limx/kggseq/). We screened for coding, non-synonymous and loss-of-function SNPs in the PPIA gene. We retained only rare variants with minor allele frequency (MAF) < 1% that were absent from the internal healthy control cohort. The gnomAD^13^ (https://gnomad.broadinstitute.org/) database was used to determine allele frequency in Non-Finnish Europeans. The whole genome sequence data is publicly available on dbGaP at phs001963.

### Molecular dynamics (MD) simulations

The structure of WT PPIA was retrieved from PDB 1CWA (ligand coordinates were removed). The mutant form was generated by swapping the lysine 76 with glutamate using UCSF Chimera ^33^. MD simulations were carried out in Gromacs 2018 ^73^ and protein topologies were generated with Charmm36m ^74^ (explicit solvent, TIP3P water). The following protocol was employed for both the WT and K76E PPIA. Each protein structure was positioned in a cubic box with 10 Å minimum distance from the walls. The system was solvated, the protein net-charge neutralized with counterions (1 Cl^-^ for the WT, 1 Na^+^ for K76E) and the final NaCl concentration was brought to 0.15 M. Then energy minimization was carried out using the steepest descent algorithm with tolerance set at 100 kJ/(mol·nm). The system was then equilibrated for 1 ns in the NVT ensemble, followed by another 1 ns equilibration in the NPT ensemble. NVT was equilibrated using the V-rescale thermostat ^75^ with reference temperature 310 K; NPT equilibration was carried out using the V-rescale thermostat (310 K) and the Parrinello-Rahman Barostat ^76^ with reference pressure 1 bar. During both equilibrations, position restraints on heavy atoms were applied with force constant 1000 kJ/(mol·nm^2^). After equilibration, position restraints were removed and a production simulation of the total length of 1 μs was done. This procedure, starting from NVT equilibration, was repeated three times for both WT and mutant PPIA, yielding a cumulative simulation time of 3 μs for each structure. Equilibrations and productions were carried out using the leap-frog integrator with a 2 fs step. The cutoff for Coulomb short-range interactions was set at 12 Å, and particle mesh Ewald was employed to treat long-range electrostatics. The cutoff for Van der Waals interactions was set at 12 Å using force-switch with radius 10 Å. RMSD and RMSF of MD trajectories were computed using Gromacs 2018; graphs were plotted using Matplotlib ^77^ in Python-3, and images of protein structures were produced in UCSF Chimera.

### Mutant PPIA cloning

The construct coding for mutant PPIA K76E was obtained by site-directed mutagenesis of the original plasmid encoding wild-type PPIA (Origene #RC203307, NM_021130) using the following forward (F) and reverse (R) primers: K76E-F, 5’-GCCATAATGGCACTGGTGGCGAGTCCATCTAT-3’, K76E-R, 5’-CAAATTTCTCCCCATAGATGGACTCGCCACCA-3’. The entire sequence of PPIA K76E was confirmed by sequencing using the following forward and reverse primers: F:5’-TTTGCAGACAAGGTCCCAAA-3’; R:5’-GTCCACAGTCAGCAATGGTG-3’.

### Cell lines, transfection and treatments

Experiments were done on HEK293 cells (ATCC CRL-1573). HEK293 cells were cultured in Dulbecco’s modified Eagle’s medium (DMEM), high glucose, GlutaMAX (Thermo Fisher), containing 10% fetal bovine serum (Thermo Fisher), 1% non-essential amino acids (Thermo Fisher) and a 1:100 dilution of penicillin/streptomycin (Thermo Fisher), at 37°C in a humidified atmosphere at 95% air and 5% CO2.

For transfection experiments, HEK293 cells were transfected with plasmids containing PPIA WT or PPIA K76E constructs, using Lipofectamine 2000 (Thermo Fisher). In detail, HEK293 cells were incubated with 0.016 μg/μL plasmid solution prepared with an Opti-MEM® Reduced-Serum Medium (Thermo Fisher), at 37°C in a CO2 incubator for 24-48 hours before testing for transgene expression.

For cycloheximide (CHX) experiments, HEK293 cells transfected with PPIA constructs were treated with 100 µg/mL of CHX for 15, 30, 45 and 60 minutes. For protein extraction, HEK293 cells transfected with PPIA constructs and/or treated with CHX were resuspended in hot buffer (1% (w/v) SDS), and DNA was fragmented using a syringe. Lysates were centrifuged at 10,000 xg for 15 min, and proteins were quantified with the BCA protein assay (Pierce).

### Immunoprecipitation

Magnetic beads coupled with sheep polyclonal antibodies anti-rabbit IgG (Dynabeads, Invitrogen) were used for co-immunoprecipitation studies. Cells were lysed in 50 mM Tris-HCl, pH 7.2, 2% CHAPS, protease inhibitor cocktail (Roche) and quantified with the BCA protein assay (Pierce). Proteins (500 µg) were diluted to 0.5 µg/µL with lysis buffer. Magnetic beads with coupled sheep antibodies anti-rabbit IgG (Dynabeads® M280; Invitrogen) were washed with 0.5% bovine serum albumin immonuglobulin-free (BSA Ig-free) in PBS to remove preservatives. Three μg of rabbit polyclonal anti-human TDP-43 primary antibody (Proteintech; RRID: AB_2200505) or rabbit polyclonal anti-human Ran primary antibody (Abcam) was incubated with 20 μL of anti-rabbit IgG-conjugated Dynabeads for 2h at 4°C in 0.1% BSA/PBS. Lysate was pre-cleared by incubation for 2h at 4°C with 30 μL of beads and incubated overnight at 4°C with primary antibody linked to the beads. After two washing steps, first with 50 mM Tris-HCl, pH 7.2, 0.3% CHAPS, 0.1% BSA Ig-free and protease inhibitor cocktail (Roche), and then with 50 mM Tris-HCl, pH 7.2, 0.3% CHAPS, and protease inhibitor cocktail (Roche), immunoprecipitated proteins were eluted by 30 μL Laemmli sample buffer, DTT 1 mM and analysed by Western blot. Controls for IP experiments were: lysate with magnetic beads linked to the secondary antibody (beads-IgGII); lysis buffer with beads-IgGII; lysis buffer with beads-IgGII and primary antibody. Immunoprecipitation experiments were repeated several times on independent sample sets, with consistent results.

---

**Supplementary Fig. 1.**
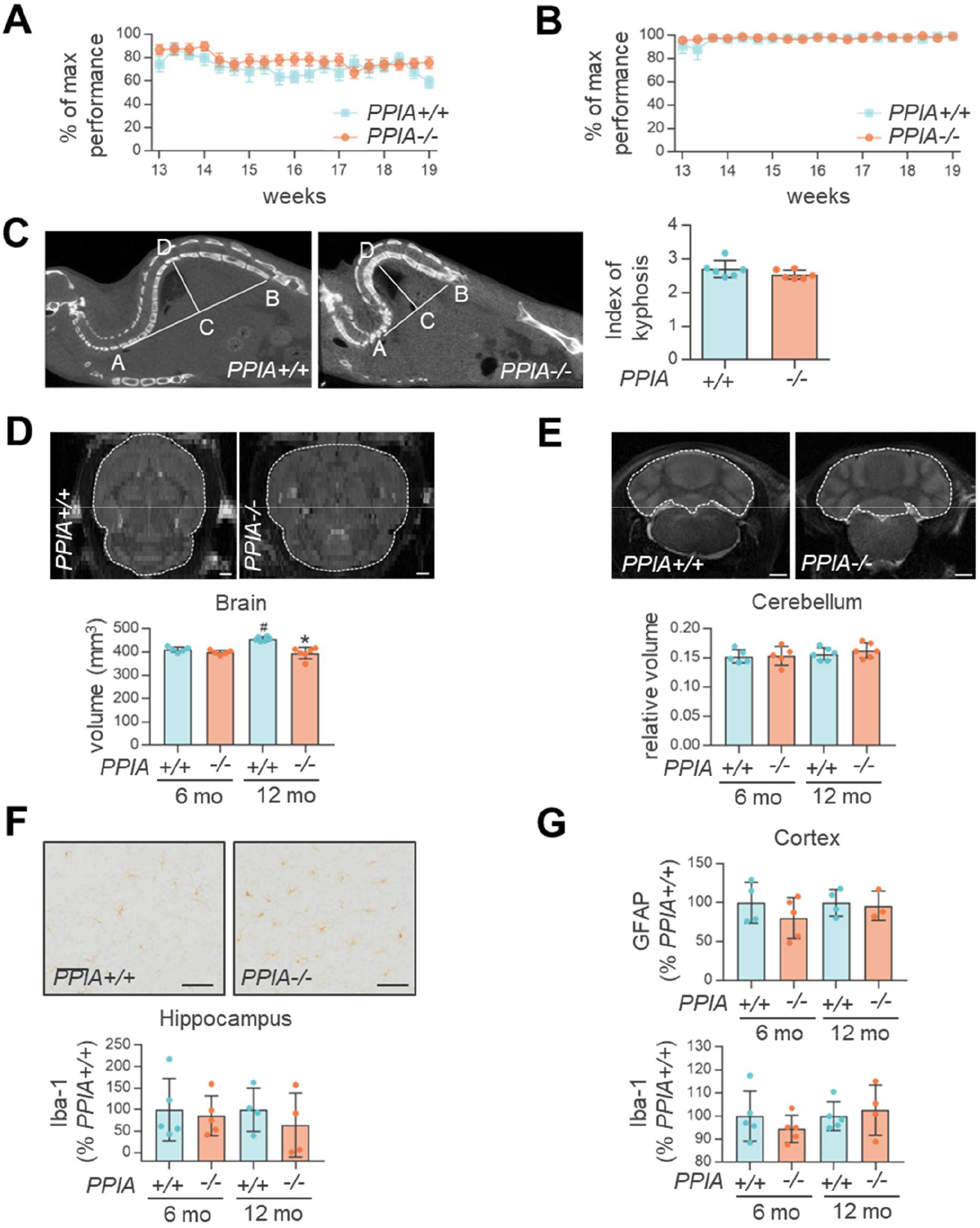
*PPIA-/-* mice show neuropathological alterations in specific brain regions. (**A-C**) Up to 20 weeks of age, *PPIA-/-* mice display no motor phenotype, but have a tendency to kyphosis at four months. *PPIA+/+* (n=10) and *PPIA-/-* (n=14) mice show a similar motor phenotype in Rotarod (**A**) and grid tests (**B**). Data (mean ± SEM) are expressed as a percentage of maximum performance and were analysed by two-way ANOVA, Bonferroni’s post hoc test. **(C)** Micro-CT analysis shows only a tendency to kyphosis in *PPIA-/-* mice compared to controls, at four months of age. Kyphosis was evaluated as the ratio between the distance from the last cervical vertebra to the last lumbar vertebra (segment AB) and the perpendicular distance to the dorsal edge of the vertebra at the greatest curvature (segment CD), as shown in the micro-CT images. Data are mean ± SD (n=6) and were analysed with Student’s t-test. (**D-E**) The volume of total brain **(D)** and cerebellum **(E)** was measured using quantitative MRI in *PPIA+/+* and -/- mice, at 6 and 12 months. Representative MRI images of *PPIA+/+* and -/- brain regions at 12 months of age are shown. The white dotted line indicates the ROI for MRI quantification. Scale bar 1 mm. The volume of the cerebellum was adjusted for total brain volume and data are expressed as relative volume. Mean ± SD (n=6 or 5 in each experimental group). *p <0.05, PPIA -/- versus *PPIA+/+* mice and ^#^p < 0.05, 6 months versus 12 months, one-way ANOVA, Tukey’s post hoc test. **(F)** Iba-1 immunostaining in hippocampus was quantified in *PPIA+/+* and -/- mice at 6 and 12 months of age. Representative Iba- 1-stained brain sections are shown. Scale bar 50 µm. Data (mean ± SD; n=5 or 4 in each experimental group) are percentages of immunoreactivity in *PPIA+/+* mice. **(G)** Dot blot analysis of GFAP and Iba-1 in cortex of *PPIA+/+* and -/- mice at 6 and 12 months. *PPIA+/+* and *PPIA-/-* mice have similar levels of GFAP (upper panel) and Iba-1 (lower panel). Immunoreactivity was normalized to protein loading. Data (mean ± SD,n=3 or 5 in each experimental group) are percentages of immunoreactivity in *PPIA+/+* mice. There was no significant difference between groups, one-way ANOVA, Tukey’s post hoc test.

**Supplementary Fig. 2.**
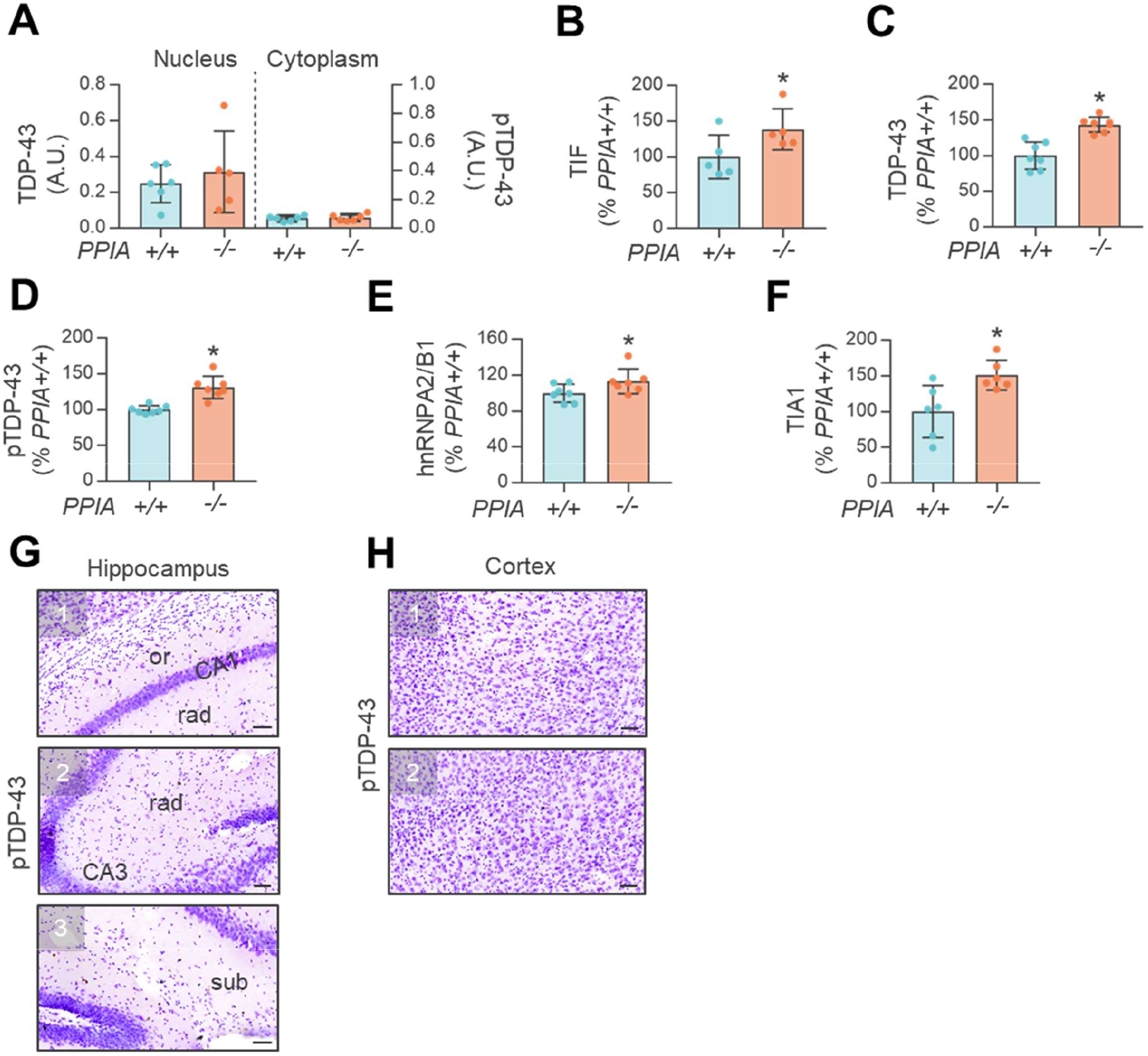
TDP-43 pathology and alterations in other RNA binding proteins in *PPIA-/-* mice. **(A)** Equal amounts of nuclear and cytoplasmic fractions from cerebellum of *PPIA+/+* and -/- mice at 6 months of age were analysed for TDP-43 or pTDP-43. Immunoreactivity was normalized to protein loading. Data (mean ± SD, n=6or 5 in each experimental group) indicates the immunoreactivity normalized to total protein loading, in arbitrary units (A.U.).and were analysed with Student’s t-test. **(B)** Analysis of the Triton-insoluble proteins (TIF) from cortex of *PPIA+/+* and -/- mice at 6 months. TIF is normalized to soluble proteins. Data (mean ± SD, n=5) are percentages of immunoreactivity in *PPIA+/+* mice. *p <0.05 Student’s t test. **(C-F)** The level of TDP-43 **(C)**, pTDP-43 **(D)**, hnRNP A2/B1**(E)**, TIA1 **(F)** in the TIF from cortex of *PPIA+/+* and -/- mice at 6 months, measured by dot blot with the specific antibodies. Immunoreactivity was normalized to protein loading. Data (mean ± SD; TDP-43, pTDP-43 and hnRNPA2/B1: n=7; TIA1: n=6are percentages of immunoreactivity in *PPIA+/+* mice. *p <0.05, Student’s t test. **(G-H)** pTDP-43 immunostaining was analysed in hippocampus **(G)** and cortex **(H)** of *PPIA+/+* mice at 12 months of age. Scale bar 50 µm.

**Supplementary Fig. 3.**
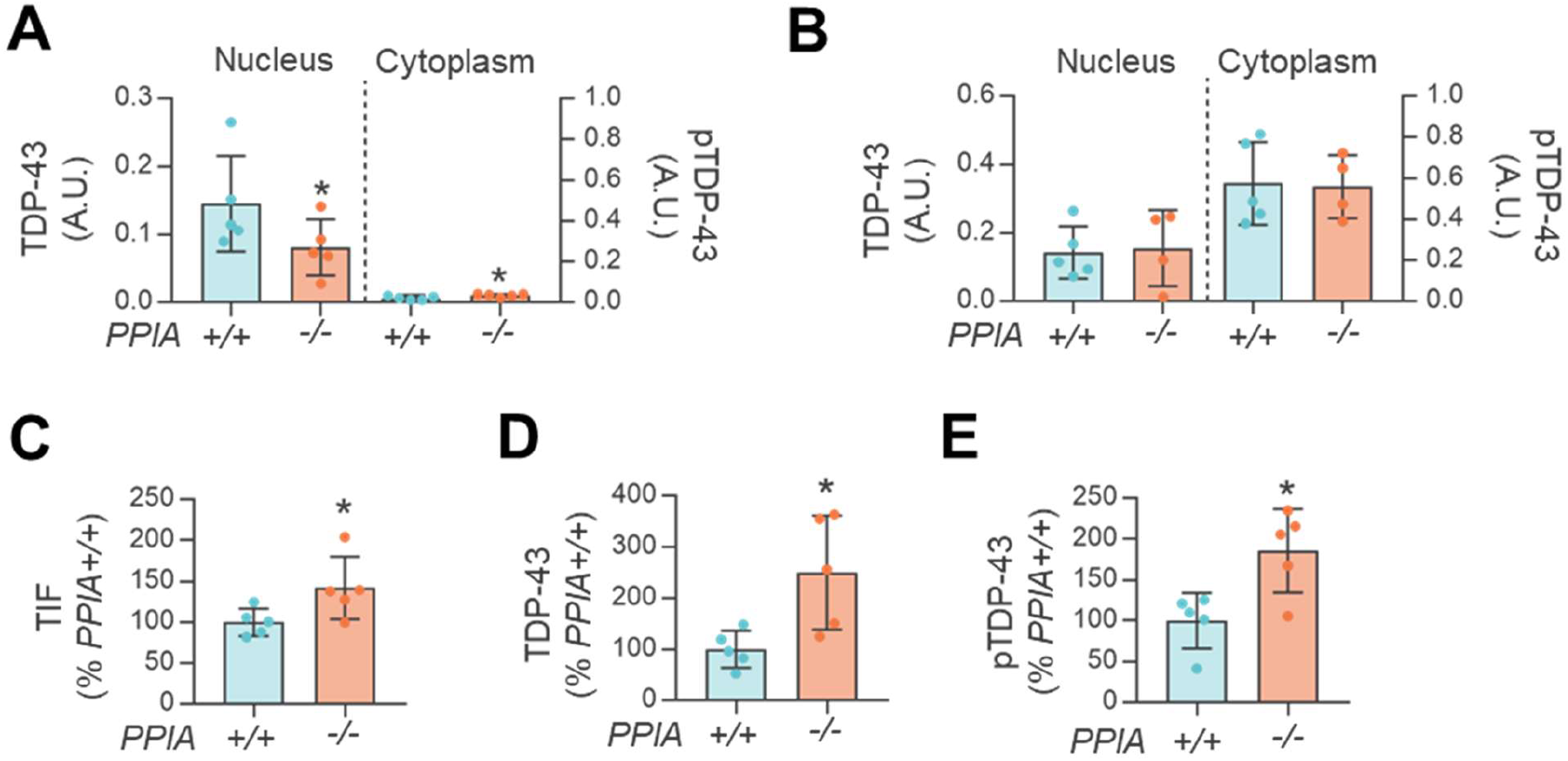
C57_*PPIA-/-* mice present TDP-43 pathology. Equal amounts of nuclear and cytoplasmic fractions from cortex **(A)** and cerebellum **(B)** of C57_*PPIA+/+* and -/- mice at six months of age were analyzed for TDP-43 or pTDP-43 by dot blot. Immunoreactivity was normalized to protein loading. **(A-B)** Data (mean ± SD, n=5 or 4 in each experimental group) indicates the immunoreactivity normalized to total protein loading, in arbitrary units (A.U.) and were analysed with Student’s t-test. **(C)** Analysis of the total TIF from cortex of C57_*PPIA+/+* and -/- mice at six months of age. Total TIF is the amount of TIF isolated from the specific tissue and is the ratio of TIF to soluble proteins. The levels of insoluble TDP-43 **(D)** and pTDP-43 **(E)** in cortex of *PPIA+/+* and -/- mice at six months of age were measured by dot blot with the specific antibodies. Immunoreactivity was normalized to protein loading. **(C-E)** Data (mean ± SD, n=5) are percentages of immunoreactivity in C57_*PPIA+/+* mice. *p< 0.05, Student’s t test.

**Supplementary Fig. 4.**
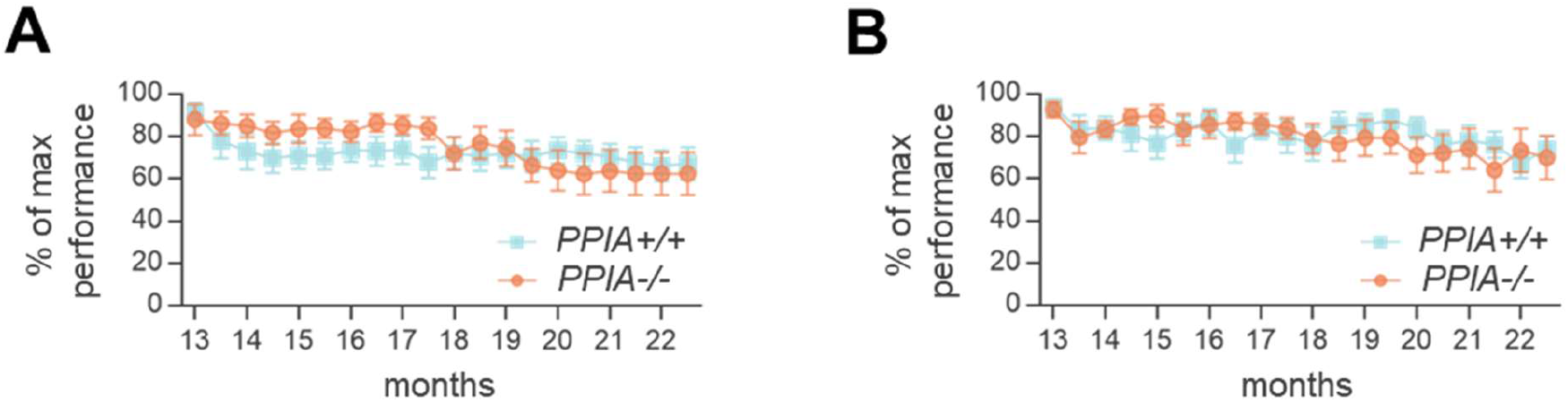
*PPIA-/-* mice present no evident motor dysfunction. Rotarod test (**A**) and grid test (**B**) show similar motor phenotypes in *PPIA+/+* (n=13) and *PPIA-/-* (n=14) mice up to 22 months. Data (mean ± SEM) are expressed as percentages of maximum performance and were analysed by two-way ANOVA and Bonferroni’s post hoc test.

**Supplementary Fig. 5.**
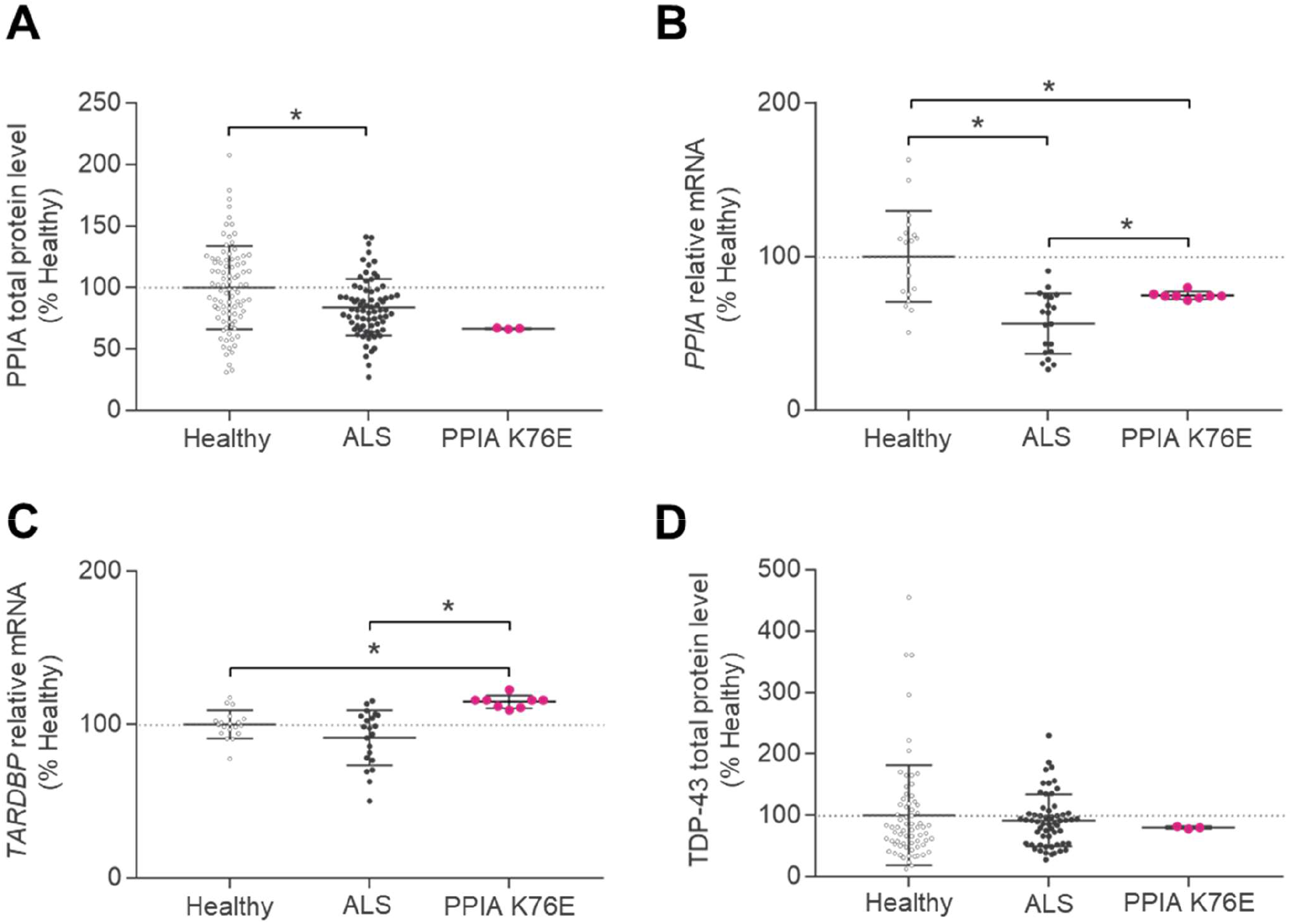
Analysis of TDP-43 and PPIA in the PPIA K76E ALS patient. (**A,B**) Analysis of PPIA protein **(A)** and transcript **(B)** levels in the PPIA K76E patient. **(A)** Total PPIA protein was analysed in PBMCs from the PPIA K76E ALS patient in comparison with ALS patients and healthy subjects of a retrospective cohort^7^ (Cohort #1, Supplementary Table1).. Scatter dot plots (mean ± SD; n=89 healthy controls, n=74 ALS patients; n=3 technical replicates of the PPIA K76E ALS patient) are percentages of healthy controls and the dotted line indicates the mean of healthy controls. *p < 0.05 versus healthy controls, Student’s t-test. (**B**) Real-time PCR for PPIA mRNA transcripts was done in PBMCs of PPIA K76E ALS patient, healthy controls and ALS patients (Cohort #2, Supplementary Table1). Data (mean ± SD; n=19 healthy subjects; n=21 ALS patients; n=8 technical replicates of the PPIA K76E ALS patient) are normalized to β-actin and expressed as percentages of healthy controls relative mRNA. The dotted line indicates the mean value of healthy controls. *p <0.05 versus healthy controls, Student’s t-test. **(C,D)** Analysis of TDP-43 transcript **(C)** and protein **(D)** levels in a PPIA K76E patient. **(C)** Real-time PCR for *TARDBP* mRNA transcripts was done in PBMCs of the PPIA K76E ALS patient, healthy controls and ALS patients (Cohort #2, Supplementary Table1). Data (mean ± SD; n=19 healthy subjects; n=21 ALS patients; n=8 technical replicates of the PPIA K76E ALS patient) are normalized to β-actin and expressed as percentages of healthy controls’ relative mRNA. The Dotted line indicates the mean of healthy controls. *p <0.05 versus healthy controls, Student’s t-test. **(D)** Total TDP-43 protein levels were analysed in PBMCs from the PPIA K76E ALS patient, ALS patients and healthy subjects of a retrospective cohort^7^ (Cohort #1, Supplementary Table1). Scatter dot plots (mean ± SD; n=69 healthy controls, n=65 ALS patients; n=3 technical replicates of the PPIA K76E ALS patient) are percentages of healthy controls and dotted line indicates the mean of healthy controls. *p <0.05 versus healthy controls, Student’s t-test.

**Supplementary Fig. 6.**
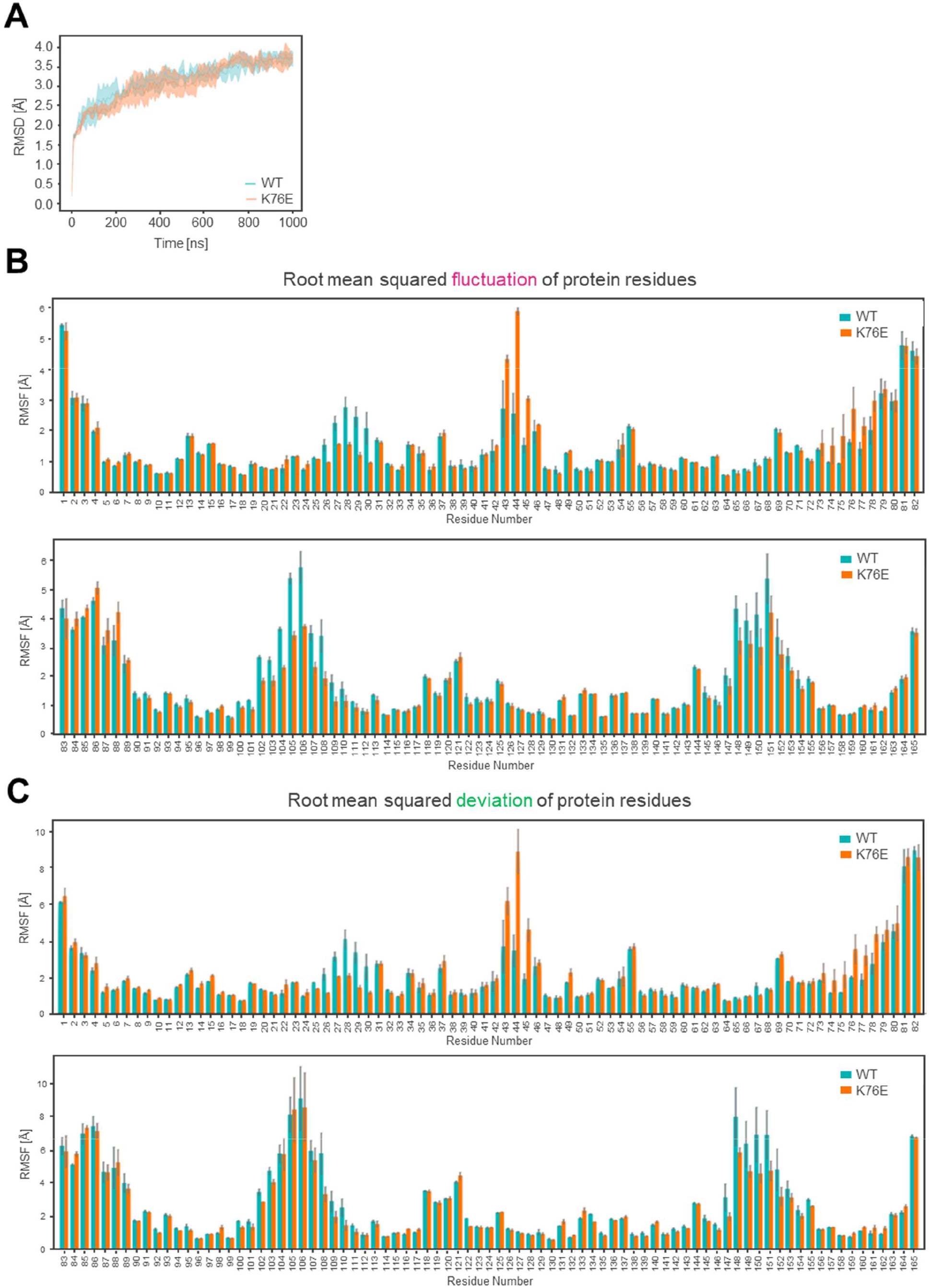
All-atom RMSD and RMSF and RMSD of protein residues computed from the MD simulations of WT and K76E PPIA. (**A**) The graph presents the all-atom RMSD of WT (blue) and K76E (orange) PPIA plotted as a function of the simulation time. Lines and filled curves represent the mean and standard error of the RMSD, respectively, computed for the three replicates of the WT and K76E protein. (**B,C**) The RMSF (**B**) and the RMSD (**C**) of the protein residues are shown. RMSD is computed using the initial frame as reference state. The WT bars are depicted in blue, and the K76E bars in orange. Vertical lines indicate standard errors.

**Supplementary Table 1.** The characteristics of the patients and controls

| Characteristics | Cohort #1 |  | Cohort #2 |  | Cohort #3 |  |
| --- | --- | --- | --- | --- | --- | --- |
|  | ALS | Healthy | ALS | Healthy | ALS | Healthy |
| No. | 79 | 89 | 21 | 19 | 51 | 20 |
| Age at sampling, years, median (range) | 65 (38-87) | 61 (37-87) | 69 (58-72) | 68 (60-72) | 67 (42-89) | 61 (53-66) |
| Sex (male/female) | 42/37 | 39/50 | 10/11 | 14/6 | 26/25 | 7/13 |
| Site of disease onset (bulbar/spinal) | 16/59 | - | 6/15 | - | 14/36 | - |
| FTD<sup>1</sup> | 5 | - | 0 | - | 1 | - |
| ALSFRS-R at sampling, median (range) | 27 (8-46) | - | 25 (10-38) | - | 33 (10-46) | - |
| ΔALSFRS-R<sup>2</sup>, median (range) | 0.9 (0.2-5.0) | - | 0.8 (0.3-1.3) | - | 0.7 (0.2-4.7) | - |
| Symptom duration<sup>3</sup>, months, median (range) | 23 (4 – 110) | - | 15 (9-101) | - | 15 (5-114) | - |

Cohort #1: PBMC samples analysed for PPIA and TDP-43 protein levels
Cohort #2: PBMC samples analysed for *PPIA* and *TARDBP* mRNA transcript
Cohort #3: Plasma samples analysed for MMP-9
<sup>1</sup>FTD: No. of patients with ALS co-morbid with FTD, according to Consensus Criteria<sup>31,32</sup>
<sup>2</sup> $\Delta$ ALSFRS-R: 48–ALSFRS-R score at sampling/time from symptom onset to sampling
<sup>3</sup>Symptom duration: Months from symptom onset to sampling

